# Sensory stimuli and cilium trafficking defects trigger the release of ciliary extracellular vesicles from multiple ciliary locations

**DOI:** 10.1101/2025.06.06.658075

**Authors:** Teresa Lobo, Guus H. Haasnoot, Aleksandra Nawrocka, Christine W. Bruggeman, Adria Razzauti, Erwin J.G. Peterman, Patrick Laurent

## Abstract

The primary cilium is a signaling organelle that extends from many cell types to detect and relay extracellular signals. Beyond its signaling role, the cilium also produces cilia-derived extracellular vesicles (cEVs), although the mechanisms underlying their biogenesis and functions remain poorly understood. We characterized the cEV biogenesis in vivo using ciliated sensory neurons of *C. elegans*. In response to sensory cues, interruption of the intraflagellar transport (IFT) -a ciliary trafficking machinery carrying cargoes along the cilium-occurs together with ciliary membrane fission, resulting in the release of cEVs. Similarly, mutants disrupting IFT and ciliary receptor trafficking also enhance cEV production. To investigate how IFT influences the rate and location of cEV biogenesis, we selected a membrane marker that spans the entire length of the ciliary membrane independently of IFT. Single-molecule tracking demonstrates that the tetraspanin TSP-6 enters and diffuses within the cilia and does it independently of IFT. Lack of receptor retrieval or receptor entry in the cilium induces membrane budding from ciliary or periciliary membranes, respectively. Prior to fission, these membrane buds get enriched in TSP-6 as well as signaling receptors. Coupling receptor buildup with their export by cEVs provides a mechanism to preserve ciliary function and to modulate ciliary signaling.

**HIGHLIGHTS:**

- The cone-shaped tetraspanin TSP-6 crosses the transition zone and moves by passive diffusion within the ciliary membrane, independently of IFT.
- The production of ciliary EVs loaded with TSP-6 increases upon acute sensory stimulation or when IFT of ciliary membrane proteins is disrupted.
- Depending on the nature of the perturbations, ciliary EVs remove excess material from the distal, proximal or periciliary segments of the cilia.

## INTRODUCTION

Cilia are microtubule-based organelles that extend from the eukaryotic cell surface to mediate sensory and motility functions^1^. Cilia are enriched with receptors and signaling machinery and are separated from the rest of the cell by a diffusion barrier, facilitating compartmentalized signaling. Key structural components common to all cilia include the transition zone (TZ) and the intraflagellar transport (IFT) complexes. Over the past decade, our understanding of the structure and function of the IFT and the TZ has advanced significantly^2^. The TZ acts as a gate-like structure at the base of the cilium and is formed by two protein complexes: the Meckel-Gruber syndrome (MKS) and the Nephronophthisis (NPHP) complexes^3^. Prior to entering cilia, the IFT-A and IFT-B subcomplexes assemble into polymeric chains known as the “IFT trains” that recruit motor proteins^4,5^. The recruitment and mobilization of these IFT trains allows for regulated bidirectional transport of ciliary cargo across the TZ and along the axonemal microtubules^6,7^.

A coordinated IFT-mediated receptor trafficking is critical for ciliary signaling. Several G-protein coupled receptors (GPCRs) associate with IFT complexes through adaptor proteins like TULP3 and the BBSome complex^8^. TULP3 facilitates the entry of the membrane-bound GPCRs across the TZ and their anterograde transport within cilia^9–12^. Conversely, the BBSome complex enables GPCR retrieval and mediates their retrograde transport and ciliary exit^13,14^. In addition, ARL13b plays a regulatory role in both the ciliary entry and exit of GPCRs^15–18^. However, whether these trafficking principles extend to other ciliary transmembrane proteins remains poorly understood.

Besides their role in signaling, cilia were observed since the 1970s to also serve as specialized sites for the formation of cilia-derived extracellular vesicles (cEVs)^19–23^. However, their biogenesis and functional significance have remained incompletely understood. As these EVs carry ciliary cargo, including membrane receptors, their shedding may serve as a mechanism to modulate ciliary signaling^21^. We previously reported that release of cEVs from *C. elegans*’ cilia alleviates the accumulations of ciliary receptors caused by receptor overexpression or impaired IFT function^24^. Previous publications in *C. elegans* highlight a diversity of EVs subtypes that can be released from distinct sites of the cilium and that contain a diversity of ciliary cargos^23,25–30^. However accurately quantifying the cEV release rates and map their budding locations in wild-type and IFT mutants has remained challenging. A universal cEV marker should uniformly localize along the whole ciliary membrane, its localization should not be directly affected by the ciliary mutations tested, it should be incorporated onto cilia-derived EVs.

The tetraspanins CD9, CD81, CD82, and CD151 are four-pass transmembrane proteins enriched in cilia in cell culture^31^. Here, we establish the tetraspanin TSP-6, the *C. elegans* ortholog of CD9 with 46% sequence similarity, as a ciliary membrane marker to explore the biogenesis of cilia-derived EVs^32^. TSP-6::wSc is an ideal cEV marker to explore the biogenesis of cEVs as it localizes along the whole ciliary membrane of several sensory neurons of *C. elegans*. It allows exploration of the effects of IFT or TZ disruptions on the cEV because TSP-6::wSc is not actively transported by IFT and spontaneously crosses the TZ. Instead, TSP-6::wSc moves within the ciliary membrane by lateral diffusion. Interestingly, we found that local properties retain and concentrate TSP-6::wSc in membrane regions prior to their outward budding into cEVs. Ciliary EV biogenesis is enhanced in mutants disrupting membrane receptor trafficking and during acute sensory signaling that transiently disrupts IFT. In these scenarios, the buildup of transmembrane proteins coincides with the sites of cilia-derived EV formation, suggesting that membrane receptor crowding may directly regulate both the location and rate of EV biogenesis.

## RESULTS

### Golgi-derived vesicles deliver TSP-6::wSc to the ciliary terminal through a kiss and run mechanism

In previous work, we reported the endogenous expression TSP-6 fused to codon-optimized mScarlet (TSP-6::wSc) in 10 of the 12 ciliated neurons of the amphid sensilla^24^. Here, we also detected endogenous expression of TSP-6::wSc in bilateral tail chemosensory neurons named PHA and PHB (Figure 1A). The bilateral PHA and PHB phasmid neurons project rod-shaped cilia through the phasmid sensilla pore made by PHso and PHsh glia. The fusion protein TSP-6::wSc was observed along the entire length of PHA and PHB cilia (Figure 1A, 1C). The simple anatomical structure of the bilateral phasmid sensilla, consisting of only PHA and PHB cilia embedded by PHsh glia and protruding through the phasmid pore into the environment, greatly facilitates the observation of TSP-6::wSc fluorescence, analysis and interpretation of the ectosome biogenesis rate and location.

**Figure 1:**
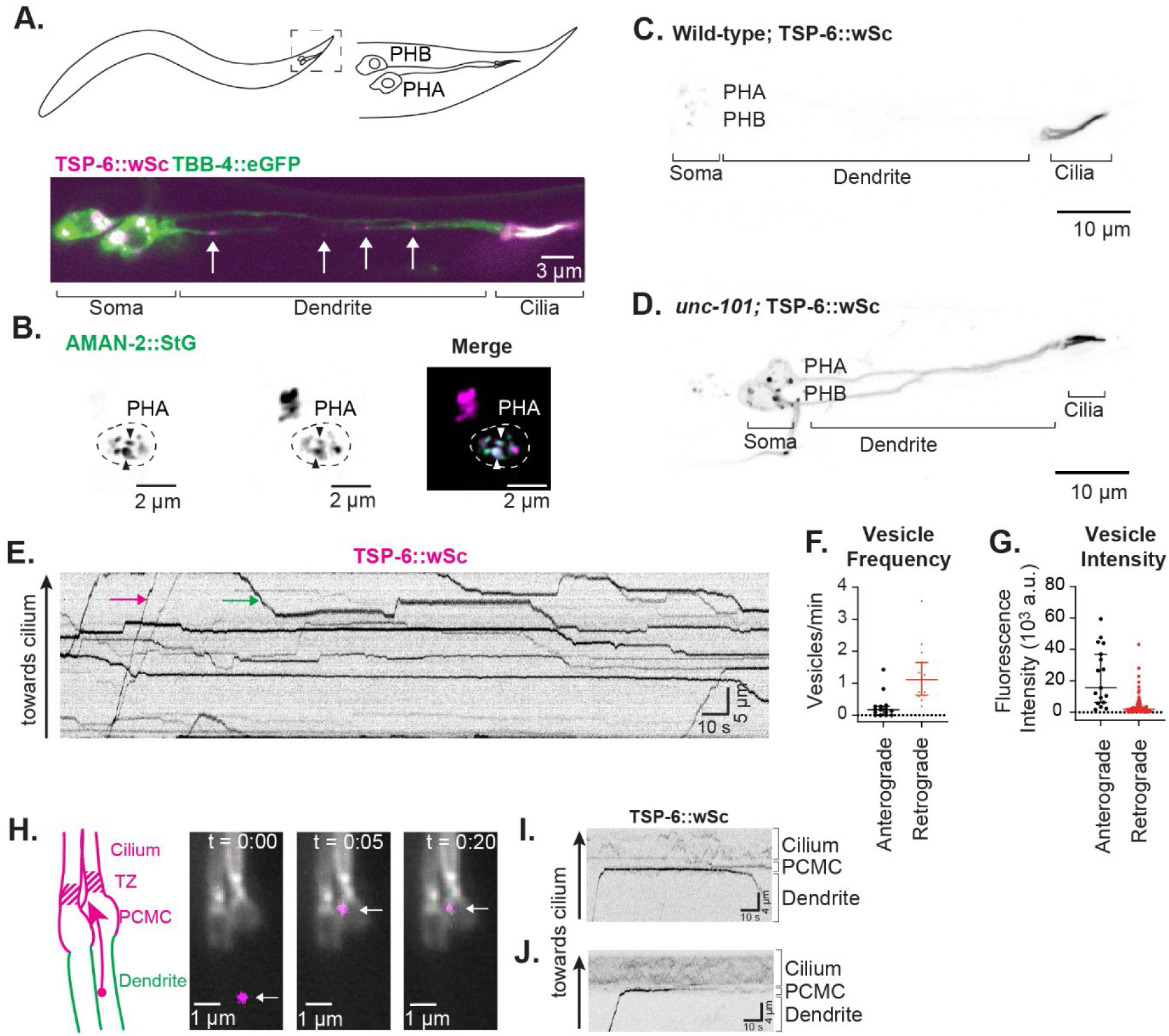
Golgi-derived vesicles deliver TSP-6::wSc to ciliated ending by kiss and run. (A) The soma of PHA and PHB neurons are in *C. elegans* tail. Their dendrites project 50um posteriorly, where their cilia enter the phasmid sensilla. Colocalization of TSP-6::wSc and TBB-4::eGFP in chemosensory neurons PHA and PHB in wild-type (N2) phasmid neurons. White arrows indicate locations of TSP-6 containing vesicles along the dendrite. (B) Colocalization of TSP-6::wSc with the Golgi marker AMAN-2::StG. Black arrows indicate puncta that colocalize. (C-D) TSP-6::wSc localized exclusively in the cilia and cell body in wild-type neurons (C), and mislocalized all throughout the membrane processes of *unc-101* neurons (D). (E) Kymograph of TSP-6::wSc taken over a length of 30 μm along the dendrite, as shown in the white bar in (A). TSP-6::wSc vesicles are highlighted moving in anterograde (magenta arrow) or retrograde (green arrow) direction toward the cilia or the soma, respectively. (F) Frequency of anterograde and retrograde TSP-6::wSc vesicle for 17 dendrites taken from 14 worms. Median anterograde frequency was 0.17 vesicle/min and retrograde was 1.11 vesicle/min. (G) Fluorescence intensity of anterograde (n = 17) and retrograde (n = 107) vesicles. The anterograde vesicles are about 8 times brighter than retrograde vesicles. However, the sum of vesicle intensity for anterograde (386761 a.u.) and for retrograde vesicles (389300 a.u.) was nearly equal. The intensities were collected from 12 worms. Error bars in vesicle frequency and vesicle intensity graphs show median with 95% CI. (H) Three consecutive frames (t=0, 5 and 20s) depict a TSP-6::wSc dendritic vesicle arriving (arrow) and immobilizing in the PCMC. Time-averaged intensity of TSP-6::wSc is shown as an overlay (greyscale); the location of the cilia,TZ, PCMC and distal dendrite are indicated in the left scheme (I) Kymograph of TSP-6::wSc vesicle arriving at PCMC and showing ‘’kiss-and-run’’ interaction. (J) Kymograph of a TSP-6::wSc vesicle arriving from the dendrite (at t=0sec) and staying immobile at the PCMC (5 to 20sec) until photobleached.

As PHA and PHB cilia localize approximately 50 µm from their soma, we were able to further characterize the biogenesis and delivery of TSP-6::wSc to their cilia. G-protein coupled receptors (GPCRs) are delivered to cilia by RAB8-positive Golgi-derived vesicles^33^. By extension, other transmembrane proteins are thought to enter Golgi-derived vesicles for delivery to cilia. Within the cell soma, the large TSP-6::wSc-positive compartment colocalized strongly with the Golgi marker AMAN-2::StayGold (StG) (Figure 1B). Some TSP-6::wSc positive vesicles colocalized with StG::RAB-8, a small GTPase involved in trafficking from the Golgi to plasma membrane (Video S1). Partial colocalization was also observed with the early endosome marker StG-RAB-5, with the recycling endosome marker StG::RAB-11 and the lysosome marker LMP-1::GFP (Videos S2-S4). No colocalization was observed with the late endosome marker StG::RAB-7 (Video S5). Within the ciliated ending, TSP-6::wSc colocalized best with the recycling endosome marker StG::RAB-11 (Video S6). Therefore, TSP-6::wSc is trafficked to the cilia in Golgi-derived vesicles, possibly via RAB-11-positive recycling endosomes. The trafficking of most ciliary transmembrane proteins from Golgi-derived vesicles to the cilium requires the AP-1 μ1 clathrin adaptor protein UNC-101, the lack of which leads to truncated cilia^33^. While TSP-6::wSc was still enriched in the cilia of *unc-101(m1)* mutants, ectopic TSP-6::wSc localization was also observed across the entire plasma membrane of the neurons, suggesting abnormal basolateral delivery of TSP-6-positive vesicles in *unc-101(m1)* (Figure 1C-D).

Kymographs of TSP-6::wSc vesicles over the entire length of the PHA/PHB dendrites revealed bidirectional vesicular trafficking within the dendrite (Figure 1E). A few bright vesicles moved towards the cilia (anterograde trafficking), with a frequency of 0.17 vesicles/min, and displayed frequent changes of direction as well as fusion and fission events (Figure 1F-G; Figures S1A-B). Many fainter vesicles moved towards the soma (retrograde trafficking) with a frequency of 1.11 vesicles/min (Figure 1F-G). These fainter retrograde tracks may correspond to vesicles resulting from TSP-6::wSc endocytosis at the periciliary membrane compartment (PCMC). The bidirectional and pausing behavior of the TSP-6::wSc vesicle traffic complicates straightforward determination of velocities. To overcome this challenge, we filtered the motility data. This filtering was performed by calculating the local anomalous diffusion coefficient *α*, the exponent that shows how the Mean Squared Displacement (MSD) scales with time. For directed motion at a constant velocity, *α* is expected to equal 2; for purely diffusive motion, *α* = 1; for subdiffusion or pausing, α < 1 (Figure S1C and S1D). We filtered the data for α ≥1.7 and determined a point-to-point velocity of 1.57 ± 0.57 μm/s (mean ± standard deviation) for anterograde particles and 0.93 ± 0.26 μm/s for retrograde particles. These mean anterograde and retrograde velocities are compatible with dynein (most likely cytoplasmic dynein 1) and kinesin motor proteins (such as kinesin-3), respectively^34,35^, suggesting that the vesicles moved towards and away from the cilia along microtubules, which are oriented with their minus-ends out in *C. elegans* dendrites (Figure S1E-F). Overall, the retrograde tracks display lower *α* values than anterograde tracks, indicating a less directed and/or more diffusive motion type (Figure S1G). Taken together, TSP-6::wSc dendritic kymographs and the point-to-point velocity analysis suggest a ‘tug-of-war’ dynamics of TSP-6::wSc vesicles, with vesicles changing direction frequently and pausing.

To investigate how TSP-6::wSc is delivered to the ciliary membrane, we photobleached the whole cilia. The delivery of bright TSP-6::wSc vesicles from the distal dendrite into the ciliated endings of the phasmid neurons could then be observed without being obscured by normally high TSP-6::wSc expression in that region (Figure 1H-J). Transient contact was observed between the PCMC membrane and TSP-6::wSc vesicles, which can remain stationary for an extended period before partly detaching or moving away (Figure 1I, Video S7-8). The absence of complete vesicle fusion with the plasma membrane suggests a kiss-and-run model, where part of the TSP-6 membrane vesicle content is diverted towards the PCMC membrane. In some cases, vesicles remained immobile while in contact with the PCMC, exhibiting a gradual decrease in intensity consistent with photobleaching, rather than the abrupt drop expected from complete fusion with the plasma membrane (Figure 1J, Video S9).

### TSP-6::wSc enters and distributes in cilia by diffusion, independent of IFT

To understand the dynamics underlying TSP-6 distribution along cilia, we performed single-molecule tracking of TSP-6::wSc in live *C. elegans* using laser-illuminated wide-field epifluorescence microscopy with a frame rate of 20 Hz (Figure 2A). Over 749 fluorescence traces collected from 14 animals were overlaid and time-integrated to map the distribution of TSP-6::wSc along the entire length of the cilia. In this analysis, data in the adjacent PHA and PHB cilia were collected in a single distribution, avoiding effects due to the partial overlap of these cilia at their distal segments. The distribution shows that TSP-6::wSc is enriched in the TZ and the distal segment of phasmid cilia (Figure 2B). Kymographs generated from the fluorescence single molecule traces were then analyzed to assess TSP-6::wSc trafficking within the cilium. Ciliary trafficking of most transmembrane proteins depends on IFT for bidirectional transport of ciliary cargo across the diffusion barrier formed by the transition zone (TZ) and along the ciliary microtubule-based axoneme although diffusive behavior is also observed^8,36^. Notably, we observed no evidence for active directional transport for TSP-6::wSc in kymographs, neither in the TZ nor in the cilium (Figures 2A, Figure S2A-B). This observation suggests that TSP-6::wSc enters strictly by diffusion across the TZ and distributes along the cilia according to its diffusion rate within the ciliary membrane, independently of IFT. From the single-molecule trajectories, we determined the diffusion coefficient of TSP-6::wSc along the cilium, which describes TSP-6::wSc rate of lateral movement within the membrane (Figure 2C). The diffusivity varied along the cilium, with the lowest value found within the TZ, the highest in the proximal cilium, and intermediate values in the distal cilium. We could not identify the mechanism responsible for this region-specific diffusivity values, which may involve differences in molecular crowding, lipid content or membrane curvature along the cilium. Using a finite difference scheme to numerically solve the one-dimensional diffusion equation, we simulated how particles spread out over time by maintaining a fixed concentration at the base and a reflective boundary at the tip (Figure 2D). This simulation uses the ciliary length, diameter, and local diffusivity of TSP-6::wSc as input parameters to predict the ciliary distribution of TSP-6::wSc. Using the measured diffusion coefficients and known ciliary dimensions, a distribution of TSP-6::wSc similar to the experimentally determined was obtained (Figure 2E).

**Figure 2:**
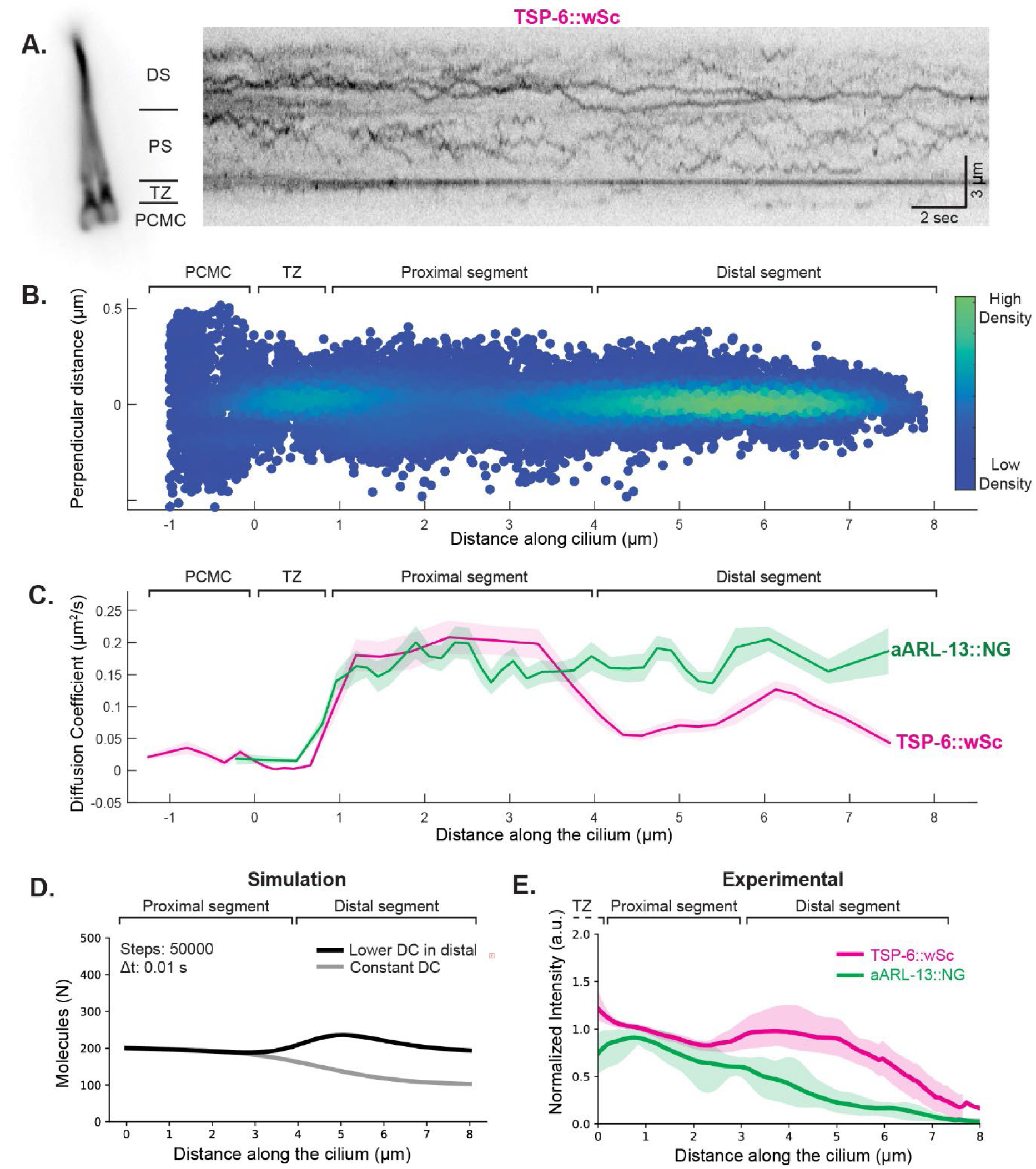
TSP-6::wSc enter and distribute in cilia by diffusion, independently of IFT. (A) Kymograph of TSP-6::wSc during photobleaching with high laser power, showing the diffusive nature of TSP-6 inside the cilium membrane and lack of IFT mediated transport. (B) Heatmap showing the time-integrated locations of tracked single-molecule TSP-6::wSc particles. The time-integrated locations were binned in 0.5 μm intervals. Within each bin, 1000 random localizations were sampled to generate this heatmap, minimizing sampling bias due to overlapping distal segments. (C) Diffusion coefficient along the length of the cilium for TSP-6::wSc and for aARL-13::NG. Average value and error are estimated using bootstrapping (see methods). (D) Distribution of molecules calculated by a diffusion simulation using a finite difference scheme to numerically solve the one-dimensional diffusion equation. The simulation ran for 50000 steps of 10 ms and accounts for the decreasing diameter of the membrane and is calculated for a constant diffusion coefficient D along the cilium or a diffusion coefficient that is 2 times lower in the distal segment than in the proximal segment. (E) Average intensity profiles taken along the spline of both cilia of TSP-6::wSc (n = 15) and aARL-13::NG (n = 4). The intensity profiles are aligned on the x-axis with their highest or lowest value in the transition zone for TSP-6::wSc and aARL-13::NG, respectively; and they are normalized on the y-axis to their maximum value in the proximal segment. Average values and error are estimated using bootstrapping (see methods).

To further validate this model, we performed the same analysis using a second diffusive membrane marker, the palmitoylated small G-protein ARL-13. This lipid-anchored protein is expected to diffuse freely and faster than a four-pass transmembrane protein of similar weight like TSP-6. We generated a truncated, GDP-locked version of ARL-13 fused to mNeonGreen, previously described to distribute along the whole cilium^37^. Single-molecule analysis of this altered ARL-13 marker (hereafter referred to aARL-13::NG) shows that it is not transported by IFT, but diffuses, with a relatively high diffusivity that appears constant along the entire cilium, unlike TSP-6::wSc which has a lower diffusion coefficient in the distal segment (Figure 2C). In the proximal segment, the diffusivity of both membrane markers is similar, suggesting that monomers of TSP-6::wSc diffuses freely within the proximal ciliary membrane, but their lateral diffusion is restrained in the distal segment, whereas aARL-13::NG diffusion is not. One-dimensional diffusion simulations accurately recapitulate the ciliary aARL-13::NG distribution (Figure 2D). Taken together, these results indicate that the ciliary distribution of TSP-6::wSc primarily reflects its local dwelling behavior in the ciliary membrane. In double transgenics, the ratio of TSP-6::wSc to aARL-13::NG increases significantly in distal cilia, suggesting a specific enrichment mechanism that retains TSP-6 in the distal ciliary segment (Figure 2E). Since TSP-6::wSc is not restricted by TZ and spans across the length of the cilia independently of IFT, we can use it to study ciliary cEV biogenesis dynamics in wild type animals and TZ or IFT mutants without risks of bias.

### Acute sensory signaling promotes ectosome biogenesis

We have previously reported that amphid cilia produce cEVs that are enriched in TSP-6::wSc^24^. Due to the densely packed anatomy of the amphid sensory organ, identifying ciliary structures and capturing cEV biogenesis in real time using conventional fluorescence microscopy was challenging. To better observe these events, we turned to the phasmid cilia in the tail of the worm. Because the physiological role of cilia is to sense the environment, we wanted to assess the potential effects of sensory activity on ectosome production. For this, we imaged PHA/PHB cilia in animals immobilized in a microfluidic device that allows acute exposure to chemicals like copper or SDS, which are known to induce calcium responses in PHA and PHB neurons^38^. Acute exposure of PHA/PHB cilia to these external cues induces a transient redistribution of IFT complexes to the ciliary base, along with shrinkage of the distal cilia and their axonemes. Subsequently, IFT resumes in the remaining proximal segment, and the cilia regrow to their full length over the course of approximately 1 hour^38^. We exposed worms expressing TSP-6::wSc to copper to observe changes happening to the ciliary membrane. Within 20 seconds of copper exposure, we observed the formation of bright puncta along the distal segment, just before the cilia appeared to fragment at their distal ends (Figure 3A-B, Video S10). The timing of this sequence suggests that distal microtubule collapse is coupled with ciliary membrane fission, resulting in the release of cEV from the ciliary tip. Because these cEV are released by budding from the apical ciliary membrane, we later refer to them as “apical ectosomes”. Kymographs indicate that these apical ectosomes remain in the phasmid pore for a prolonged period after the ciliary collapse. Within 20 second of copper exposure, kymographs of the homodimeric kinesin-2 OSM-3::mCherry show that IFT is transiently halted. During IFT interuption, the OSM-3::mCherry localized in the distal segment of the cilia enter apical ectosomes (Figure 3C). Ultimately, apical ectosomes are ejected through the phasmid pore into the environment. This was confirmed by the release of OSM-3::mCherry-positive apical ectosomes into the medium (Figure 3C, Video S11). In addition to TSP-6::wSc and OSM-3::mCherry we observed that the apical ectosomes also contained the TRPV channel OCR-2, the IFT-B subunit OSM-6::eGFP and the dynein-2 subunit XBX-1::eGFP (Figure S3A-B). In addition to the formation of apical ectosomes, we also occasionally observed the release of cEVs from the PCMC after the collapse of the distal segment induced by the chemical stimulus (Figure 3D). These cEVs, referred to as basal ectosomes hereafter, appeared released by outward budding from the PCMC and were subsequently captured by the supporting glial cell PHsh (Figure 3D, Video S12 and see below). Taken together, these results show that acute ciliary sensory stimulation can drive the budding of ectosomes from the ciliary tip. Under these conditions, different molecular components present at the ciliary distal segment can be packed and released in these apical ectosomes.

**Figure 3:**
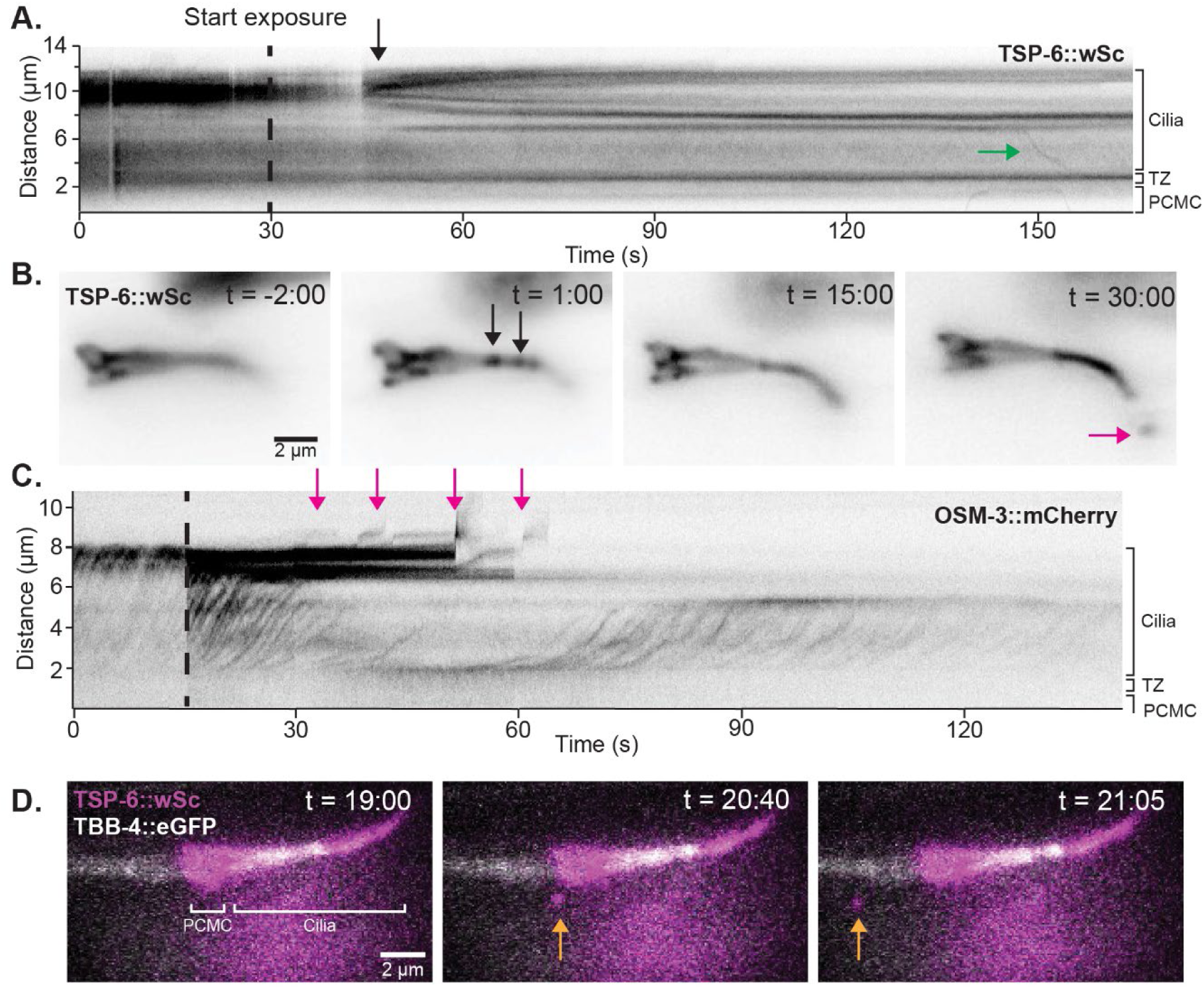
Acute sensory signaling promote ectosome budding. (A) Kymograph of TSP-6::wSc after exposure to 20 mM CuSO_4_. The start of the exposure is indicated with the black dashed line. The ciliary membrane breaks in two ∼15 sec after cues delivery (black arrow). The green arrow indicates TSP-6::wSc molecules being actively transported together in retrograde direction at an average velocity of 0.66 µm/s. (B) Time-average fluorescence intensity images of TSP-6::wSc before, during and after exposure to 20 mM CuSO_4_. TSP-6::wSc puncta -likely where membrane fission occurs-appears within 1 minute after exposure to CuSO_4_. Within 30 seconds after exposure to CuSO_4_ the TSP-6::wSc puncta disappear and an apical ectosome (magenta arrow) is ejected (D) Kymograph of OSM-3::mCherry after exposure to 0.1% SDS, dashed line indicates the onset of exposure, magenta arrows indicate the pinching and release of apical ectosomes to the environment and. (D) Subsequent time-average fluorescence intensity images of TSP-6::wSc, during basal ectosome release.

### Ciliary transmembrane protein trafficking controls the location of ectosome biogenesis

We previously reported that ectosomes can alleviate the accumulation of ciliary receptors caused by impaired IFT function^24^. We hypothesized that this could be due to an increased rate of ectosome biogenesis (ectocytosis), a shift in the ectosome budding sites, or both. To test this, we developed an assay to compare the rate of ectosome production between wild-type controls and ciliary mutant of interest (Figure 4A). We determined the rate of apical ectosome production from phasmid neurons by counting TSP-6::wSc apical ectosomes near the tail of hermaphrodite animals one hour after mounting (Figure 4B). In addition, we quantified the rate of basal ectosome production by inferring the fluorescence intensity of TSP-6::wSc basal ectosomes accumulating within the PHsh soma following their capture close to the cilia base (Figure 4C). Many GPCRs mediating phasmid sensory responses cross the TZ via anterograde IFT in a TUB-1-dependent manner and exit the cilia via retrograde IFT in a BBSome-dependent manner^9,39,40^. Therefore, we reasoned that local accumulation of GPCRs and other transmembrane proteins due to impaired entry or exit would influence ectosome budding. To test this, we measured ectosome production in *tub-1* and *bbs-8*, the latter being one of the subunits of the BBSome. We noted that TSP-6::wSc did not require *tub-1* or *bbs-8* for its ciliary localization, as expected since it enters the cilium by diffusion independent of IFT (Figure 4D). As predicted, we observed increased basal ectocytosis in *tub-1* and increased apical ectocytosis in *bbs-8* mutants (Figure 4E-F). Increased basal ectocytosis in *tub-1* correlated with PCMC expansion and ciliary shortening, whereas increased apical ectocytosis in *bbs-8* correlated with PCMC shrinkage and cilium extension (Figure 4E-H). To better pinpoint the location of ectosome budding along *bbs-8* cilia, we imaged TSP-6::wSc together with the axoneme-enriched tubulin marker TBB-4::GFP in dual-color experiments using super-resolution microscopy. We observed membrane buds enriched with TSP-6::wSc appearing along the cilia of *bbs-8* from its proximal to its distal end (Figure 4I-J). Interestingly, TSP-6::wSc appeared to be enriched in these membrane buds prior to ectosome release, suggesting a mechanism in which TSP-6 is enriched in developing ectosomes before budding, such as a preference for highly curved membranes or for local lipids. Localized buildup of receptors is observed in the cilia of *bbs-8* ^13,14,40,41^, which may sort them to the ectosome biogenesis site, as observed for GPCRs at the cilia tip ^42^. The TRPV receptor OCR-2::GFP was incorporated into apical ectosomes following acute sensory stimulation (Figure S3B). We observed OCR-2::GFP together with TSP-6::wSc in the membrane buds along the *bbs-8* cilia, suggesting coordinated sorting of OCR-2::GFP and TSP-6::wSc into ectosomes (Figure 4K).

**Figure 4.**
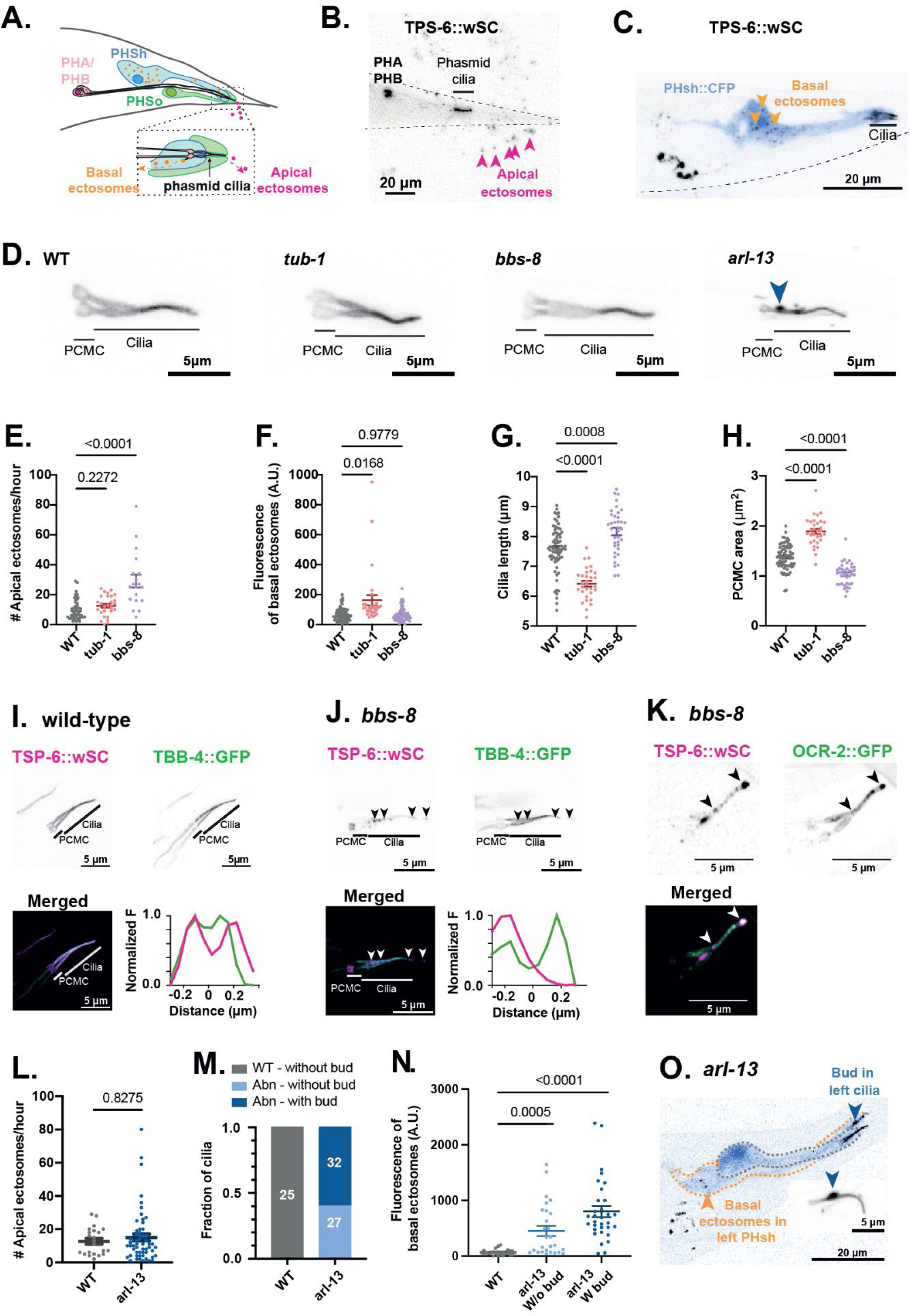
Ciliary transmembrane protein trafficking controls the location of ectocytosis. (A) PHA and PHB neurons shed ectosomes from the cilia base and from the cilia tip. The basal ectosomes released from the cilia base (orange) are captured and accumulate the soma of the PHsh glia that support PHA and PHB cilia (blue). The apical ectosomes (magenta) released from the cilia tip are released into the external environment. (B) The release of apical ectosome is quantified by counting the TSP-6::wSc positive ectosomes surrounding the tail (magenta arrowheads) one hour after mounting. (C) The release of basal ectosome (orange arrowheads) is quantified by determining TSP-6::wSc fluorescence signal within the PHsh soma. (D) The PHA and PHB cilia labelled with TSP-6::wSc exhibits different morphologies in *tub-1, bbs-8* and *arl-13* mutants. Compared to N2 controls, *tub-1* mutants show a larger PCMC and shorter cilia; *bbs-8* mutants exhibit a smaller PCMC and longer cilia; *arl-13* mutants display abnormal blebbing in the proximal cilia (blue arrowhead). (E) Apical ectosome production takes place in wild-type (N=46), *tub-1*(N=28) and *bbs-8* mutants (N=19). It is increased in bbs-8 mutants compared to wild-type controls. Kruskal–Wallis ANOVA was performed followed by Dunn’s post-hoc test for multiple comparisons. (F) Basal ectosome production takes place in wild-type (N=67), *tub-1*(N=31) and *bbs-8* mutants (N=53). It is increased in *tub-1* compared to wild-type controls. Brown–Forsythe ANOVA was performed followed by Dunnett’s T3 post-hoc test to correct for multiple comparisons. (G) Compared to wild-type controls (N=73); *tub-1*(N=31) presents shorter cilia whereas *bbs-8* (N=37) presents longer cilia. Brown–Forsythe ANOVA was performed followed by Dunnett’s T3 post-hoc test to correct for multiple comparisons. (H) Compared to wild-type controls (N=74); the PCMCs area are enlarged in *tub-1* (N=31) and reduced in *bbs-8* (N=39). Brown–Forsythe ANOVA was performed followed by Dunnett’s T3 post-hoc test to correct for multiple comparisons. (I) The axonemes microtubules are observed within the cilia of wild-type using the tubulin marker TBB-4::GFP. The fluorescence along a cross-section made at the level of proximal cilium is displayed. The TSP-6::wSc labelling appears outside the TBB-4::GFP-labeled axoneme, approximately 0,2 um from the cilium center. (J) Membrane buds enriched with TSP-6::wSc are observed alongside the cilia of *bbs-8* (arrowheads). The fluorescence along a cross-section made at the level of proximal cilium is displayed. A strong unilateral TSP-6::wSc labelling appears outside the TBB-4::GFP-labeled axoneme, approximately 0,2 um from the cilium center. (K) In *bbs-8* mutants, OCR-2-GFP appears within the membrane buds containing TSP-6::wSc (arrowheads) located at cilia membrane. (L) Apical ectosome release is potentiated in *arl-13* mutants (N=31) compared to wild-type controls (N=73). Unpaired Mann-Whitney test. (M) Abnormal (abn) cilia morphologies are observed in *arl-13* mutants: 61% display membrane budding along the proximal cilia and 39% do not display obvious budding but yet display abnormal shape (N=20 for both genotypes). (N) The *arl-13* cilia displaying membrane buds (N=32) export more basal ectosomes toward ipsilateral PHsh when compared to *arl-13* without buds (N=27) and wild-type (N=25). Brown–Forsythe ANOVA was performed followed by Dunnett’s T3 post-hoc test to correct for multiple comparisons. (O) Membrane budding in the proximal region of the left cilia in an *arl-13* mutant is associated with basal ectosome accumulation in the left PHsh, whereas no ectocytosis events are observed in the right cilia and right PHsh.

ARL13b is a small GTPase that is essential for the regulation of ciliary PI(4,5)P_2_, BBSome function and the retrograde trafficking of certain receptors^15–17,37,43^. As a result, membrane receptors accumulate in the proximal cilia of *arl-13* mutants^9,17^. In the *arl-13* mutant, we observed increased apical and basal ectosome release together (Figure 4L-O). Interestingly, 61% of *arl-13* mutants exhibited membrane buds protruding from the proximal ciliary segment; the other 39% had either a wild-type shape or abnormally shaped cilia but no evidence of ciliary budding (Figure 4M). The appearance of proximal membrane buds correlated with increased ectosome export to the ipsilateral PHsh glia (Figure 4N). When buds appeared unilaterally in the left PHA and PHB cilia, we observed ectosome-derived signal only in the left PHsh glia (Figure 4O). Therefore, buds in the proximal cilia -where receptors and TSP-6::wSc accumulate in *arl-13*-can export ectosomes to ipsilateral PHsh glia. Consequently, a close apposition of the glial membrane with the ciliary membrane is not required for ectosomes to be captured by the supporting glia. Time-lapse imaging confirmed the budding of ectosomes from the proximal cilia in *arl-13* mutants over <1 minute (Video S13). Taken together, the results in *tub-1*, *bbs-8* and *arl-13* mutants suggest TSP-6::wSc enriched ectosomes are produced at the locations where there is receptor buildup, suggesting that these receptor buildups are seeding ectosome production.

### Ciliary signaling controls the location of ectosome release

Given that mislocalized receptors causes disruption of ciliary signaling, we wondered whether long-term modulation of sensory responses can regulate the biogenesis of ectosomes. In *C. elegans,* the cGMP-dependent protein kinase EGL-4/PKG pathway reduces sensory responses^44–46^. Hyper or hypoactive sensory responses are observed in *egl-4* loss-of-function (*lf*) and gain-of-function (*gf*) alleles, respectively^47–49^. We used these mutants to assess how modulation of sensory responses impact ectosome release. We observed an increased rate of apical ectosome release in *egl-4(lf)* animals, in which sensory responses are enhanced (Figure 5A). We observed a slight increase in basal ectosome release in *egl-4(gf),* in which sensory responses are reduced (Figure 5B). PHA and PHB responses to copper and SDS depend in part on the TRPV channel OCR-2^44–46^. In *ocr-2* mutants, the rate of apical and basal ectosome production was unchanged (Figure 5C-D). Therefore, long-term enhancement of ciliary signaling weakly increases the rate of apical ectosome release, while reduced ciliary signaling slightly increases basal ectosome release.

**Figure 5.**
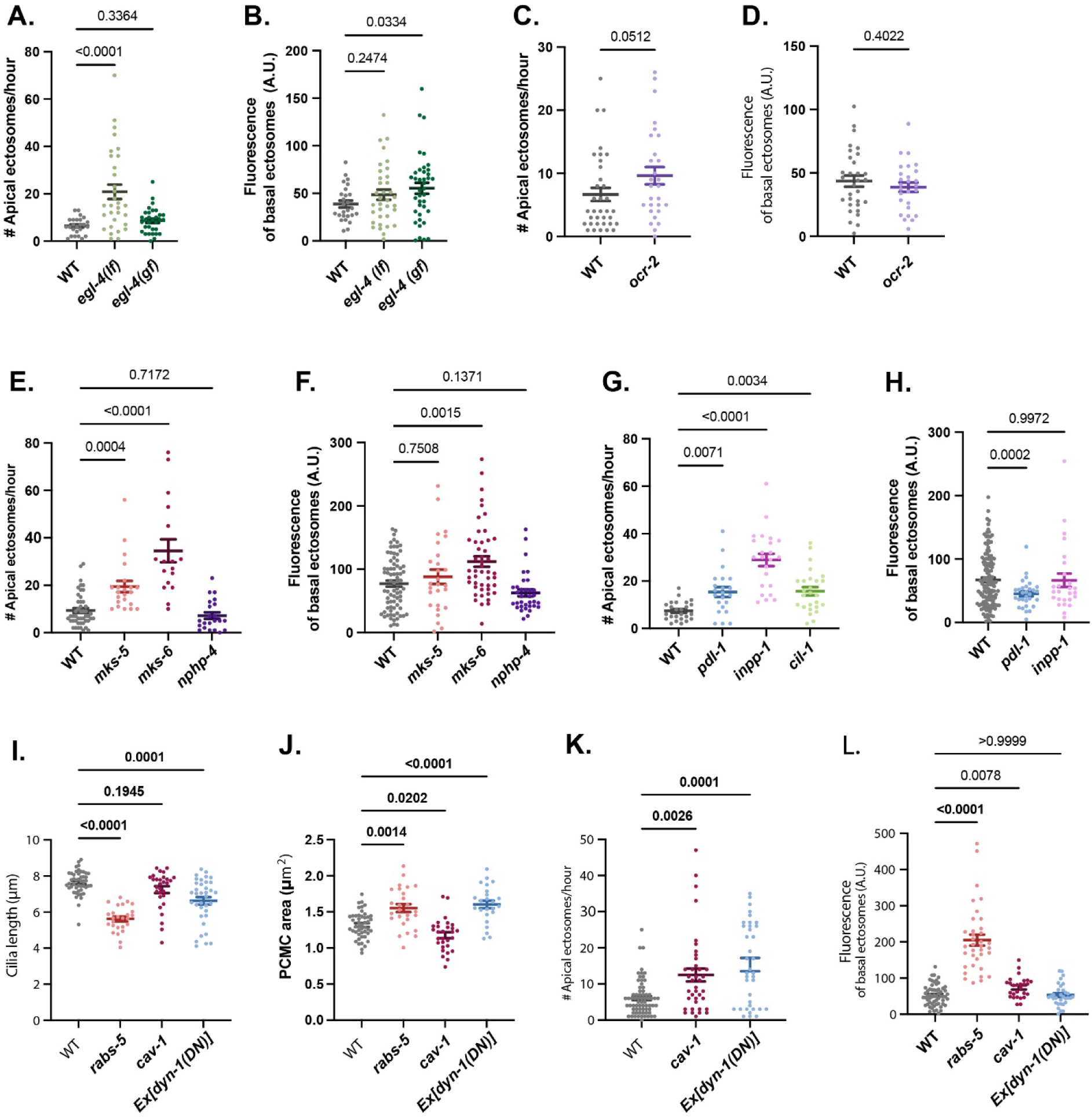
Ciliary signaling, phospholipid distribution and endocytic activity control the location of ectosome release. (A) Compared to controls (N=25), the apical ectosome release is increased in *egl-4(lf)* (N=35), but not in *egl-4(gf)* mutants (N=39). Kruskal–Wallis ANOVA was performed followed by Dunn’s post-hoc test for multiple comparisons. (B) Compared to controls (N=26), the basal ectosome release is not modified in *egl-4(lf)* (N=35) and *egl-4(gf)* mutants (N=39). Brown–Forsythe ANOVA was performed followed by Dunnett’s T3 post-hoc test to correct for multiple comparisons. (C) Apical ectosome release is not modified in *ocr-2* mutants (N=28) compared to controls (N=35). Unpaired Mann-Whitney test. (D) Basal ectosome release is not modified in *ocr-2* mutants (N=27) compared to controls (N=31). Unpaired t test with Welch’s correction. (E) Compared to controls (N=47), apical ectosome release is potentiated in *mks-5* (N=22) and *mks-6* (N=17), but not in *nphp-4* mutants (N=22). Kruskal–Wallis ANOVA was performed followed by Dunn’s post-hoc test for multiple comparisons. (F) Basal ectosome release is observed in *mks-5* (N=27), *mks-6* (N=47) and *nphp-4* mutants (N=34). It is increased in *mks-6* mutants compared to controls (N=76). Brown–Forsythe ANOVA was performed followed by Dunnett’s T3 post-hoc test to correct for multiple comparisons. (G) Apical ectocytosis is increased in *pdl-1* (N=22), *inpp-1* (N=23) and *cil-1* mutants (N=26) compared to controls (N=25). Kruskal–Wallis ANOVA was performed followed by Dunn’s post-hoc test for multiple comparisons. (H) Basal ectosome release is not modified in *pdl-1*(N=30) and *inpp-1* mutants (N=26) compared to controls (N=133). Brown–Forsythe ANOVA was performed followed by Dunnett’s T3 post-hoc test to correct for multiple comparisons. (I) Compared to wild-type controls (N=47), *rabs-5* mutants (N=26) display strongly shortened cilia; whereas *cav-1* mutants (N=29), and neuronal expression of DYN-1(K46A) (N=36) result in slightly shortened cilia. Brown–Forsythe ANOVA was performed followed by Dunnett’s T3 post-hoc test to correct for multiple comparisons. (J) Compared to wild-type controls (N=47), the PCMC area is enlarged in *rabs-5* mutants (N=26) and in animals expressing DYN-1(K46A) (N=24) in PHA and PHB neurons but reduced in *cav-1* mutants (N=29). Brown–Forsythe ANOVA was performed followed by Dunnett’s T3 post-hoc test to correct for multiple comparisons. (K) Compared to wild-type controls (N=69), the apical ectosome release is increased in *cav-1* mutants (N=38) and in animals expressing a neuronal DYN-1(K46A) construct (N=37). Kruskal–Wallis ANOVA was performed followed by Dunn’s post-hoc test for multiple comparisons. (L) Compared to wild-type controls (N=59), the basal ectosome release is increased in *rabs-5* (N=37) and in *cav-1* mutants (N=27) but not in animals expressing DYN-1(K46A) (N=32). Brown– Forsythe ANOVA was performed followed by Dunnett’s T3 post-hoc test to correct for multiple comparisons.

### Ciliary phosphatidylinositol 4,5-bisphosphate PI(4,5)P2 promotes apical ectocytosis

Together with BBS-8, TUB-1 and ARL-13, the TZ diffusion barrier regulates ciliary entry and retention of transmembrane proteins and lipids such as PI(4,5)P_2_^50^. The TZ is formed by 2 complementary modules: the MKS and NPHP modules^3,51^. Although both are involved in ciliary gating, loss of the MKS or the NPHP genes differentially affects ciliary morphology and composition^51–53^. In *nphp-4* mutant animals, lacking a key subunit of the NPHP module, we did not observe significant changes in ectosome production. In contrast, in *mks-5* or *mks-6* mutants, which lack key subunits of the MKS complex, we observed increased apical ectosome release rates (Figure 5E-F). Live imaging of TSP-6::wSc in *mks-6* mutants revealed a sequence of events for apical ectocytosis involving filopodia-like extensions from the ciliary tip membrane, followed by pinching, ectosome release, and retraction of the remaining protruding membrane (Video S14). Previous publications suggest that *mks-5*, *tub-1* and *arl-13* result in abnormal accumulation of PI(4,5)P_2_ in the cilium proper^12,43,53,54^. Ciliary PI(4,5)P_2_ is strictly regulated: it is kept enriched in periciliary membrane compartment (PCMC) by the PIP kinase PPK-1 but excluded from the cilium by TZ gating properties and by the cilia-resident PI(4,5)P_2_-degrading enzyme INPP5E (inositol polyphosphate 5-phosphatase)^12,55–57^. The prenyl-binding protein PDE6D - known as PDL-1 in *C. elegans* – helps to relocate the INPP5E enzyme from the PCMC to the cilium, across the transition zone (TZ)^43,58,59^. To assess the potential effects of increased ciliary PI(4,5)P_2_ on ciliary morphology and ectosome release, we analyzed deletion mutants for *pdl-1* and mutants for the two INPP5E orthologs in *C. elegans*: *inpp-1* and *cil-1.* All three mutants exhibited an increased rate of apical ectosome release (Figure 5G). Basal ectosome export was unaffected in *inpp-1* and decreased in *pdl-1* in which INPP-1 and CIL-1 accumulate in PCMC, likely reducing PI(4,5)P_2_ locally (Figure 5H). Ectosome release or cilia decapitation were previously observed when PI(4,5)P_2_ increases in cilium^56,60^. These results show that increased ciliary PI(4,5)P_2_ levels promote apical ectosome release.

### Endocytosis and endosome recycling regulate PCMC size and ectosome release

The ciliary distribution of PI(4,5)P_2_ and other phosphoinositides broadly affects cilium biology including endocytosis, endosome recycling, cilia entry of transmembrane protein, cilia assembly and stability^55,61^. We explored the effect of the endocytic and endosome recycling activity on cilia morphology and ectocytosis. *rabs-5* mutants disrupt both endocytic activity and endosome recycling at the PCMC, leading to the accumulation of vesicles in the PCMC^62,63^. In *rabs-5,* we observed a PCMC expansion, a reduced cilium length and an increased production of basal ectosomes (Figure 5I-J and L). Caveolin is suggested to contribute to endocytic activity at the PCMC^62–64^. In the caveolin mutant *cav-1*, we observed increased apical ectosome release, a slight increase in basal ectosome release and a modest decrease in PCMC area (Figure 5I-L). The overexpression of a dominant-negative dynamin DYN-1(K46A) has been shown to reduce clathrin-mediated endocytosis^65^. Expression of DYN-1(K46A) in PHA and PHB neurons induced PCMC expansion and increased apical ectosome release, while slightly reducing cilium length (Figure 5I-L). Combined, these results suggest that endocytic activity and endosome recycling control the PCMC size and the location of ectocytosis. Although the mechanism of action remains unclear, these results suggest that the tightly controlled import and removal of ciliary membrane proteins prevents ectosome production, and that changes in the endocytic and recycling equilibrium can promote either apical or basal release of ectosomes.

### Cilia truncation supports basal ectocytosis through PCMC expansion and branching

We have shown here that during acute sensory signaling, ectosome release carries not only the transmembrane proteins OCR-2 and TSP-6, but also the IFT-B component OSM-6 and the motor proteins XBX-1 and OSM-3 (Figure S3A-B). We used mutants of IFT-A and IFT-B complexes to explore whether the local accumulation of IFT components and motor proteins actively promotes ectosome formation or whether they passively enter ectosomes. Cilia shortening occurs in the absence of IFT-A and IFT-B complexes (Figure 6A). However, the underlying causes are different: IFT-A mutations (*che-11*, *ift-74*) prevent retrograde IFT, leading to the accumulation of IFT-B complexes at the distal tips, whereas mutations in the IFT-B complex (*osm-6*) prevents anterograde IFT, leading to the accumulation of IFT-A complexes at the ciliary base^66,67^. Despite these differences, we observed that all these mutants exhibit, similar PCMC expansion, a strongly increased rate of basal ectosome production, whereas apical ectosome production remained unchanged (Figure 6B-D). In the absence of the homodimeric kinesin-2 motor protein OSM-3, IFT is halted in the distal ciliary segment, resulting in its truncation and phenotypes similar to *che-11, ift-74* and *osm-6* regarding the PCMC area, ectosome production and cilia morphology (Figure 6A-E).

**Figure 6.**
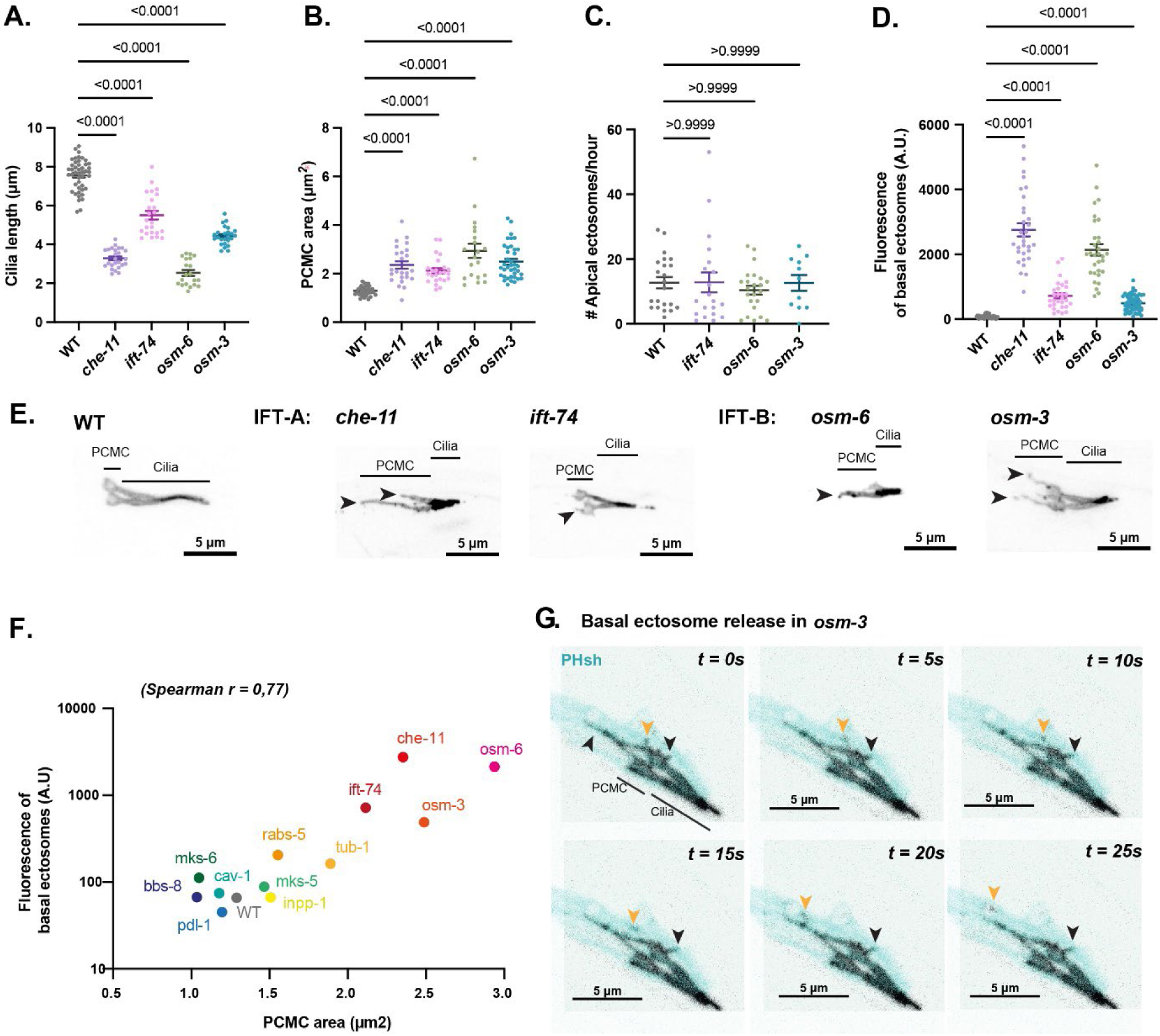
Cilia truncations support basal ectocytosis through PCMC expansion and branching. (A) The length of cilia proper is reduced in *che-11* (N=24), *ift-74* (N=24), *osm-6* (N=20) and *osm-3* mutants (N=38) compared to controls (N=34). Brown–Forsythe ANOVA was performed followed by Dunnett’s T3 post-hoc test to correct for multiple comparisons. (B) The PCMC compartments are enlarged in *che-11* (N=24), *ift-74* (N=24), *osm-6* (N=20) and *osm-3* mutants (N=38) compared to controls (N=34). Brown–Forsythe ANOVA was performed followed by Dunnett’s T3 post-hoc test to correct for multiple comparisons. (C) Apical ectosome release is not modified in *ift-74* (N=29), *osm-6* (N=23) and *osm-3* mutants (N=11) compared to controls (N=22). Kruskal–Wallis ANOVA was performed followed by Dunn’s post-hoc test for multiple comparisons. (D) Basal ectosome release is increased in *che-11* (N=31), *ift-74* (N=29), *osm-6* (N=31) and *osm-3* mutants (N=57) compared to controls (N=54). Brown–Forsythe ANOVA was performed followed by Dunnett’s T3 post-hoc test to correct for multiple comparisons. (E) PHA & PHB cilia were visualized using TSP-6::wSc. In *che-11, ift-74, osm-6* and *osm-3* mutants, variable shortening of the cilia proper are observed together with expansion and branching of the PCMCs. (F) Spearman’ rank correlation coefficient suggests a strong correlation between the PCMC area and the average production of basal ectosome (log^10^ of PHsh fluorescence). The logarithmic relationship may be explained because PHsh fluorescence result from accumulations of ectosomes over the entire life of the animals. (G) In *osm-3* mutants, dynamic branches originating from the PCMC (black arrowheads) grow and retract, these branches serve as preferential basal ectosome shedding sites (orange arrowheads).

Compared to *tub-1*, which only disturbs transmembrane ciliary protein trafficking, complete IFT disruption induces a very high rate of basal ectosome release. This enhanced basal ectosome release correlated well with increased PCMC area (Figure 6F). In addition to a PCMC expansion, we observed the formation of PCMC branches, which were also embedded in the surrounding PHsh glia (Figure 6E and 6G). To observe the process of basal ectosome release, we performed time-lapse imaging of *osm-3* mutants tracking the TSP-6::wSc signal in animals expressing a glial reporter. We observed that the PCMC is highly dynamic in *osm-3* mutants, with branches expanding and retracting within ∼4 minutes (Figure 6G; Video S15). Basal ectosomes budded outward from both PCMC branches and from the main PCMC membrane, after which they were captured by PHsh glia and transported retrogradely towards the cell soma. We observed the release of approximately one basal ectosome per minute (Figure 6F; Video S16). Although we did not observe apical ectosome release during live imaging of *osm-3* mutants, we observed apical ectosome released in the medium and surrounding the tail of *osm-3* animals. To better visualize their cilia architecture, we used the axoneme-enriched beta-tubulin marker TBB-4::GFP. At the tip of the *osm-3* cilia, we observed TSP-6::wSc membrane budding distal to the axoneme ending, likely prior to apical ectocytosis (Figure 7A).

**Figure 7.**
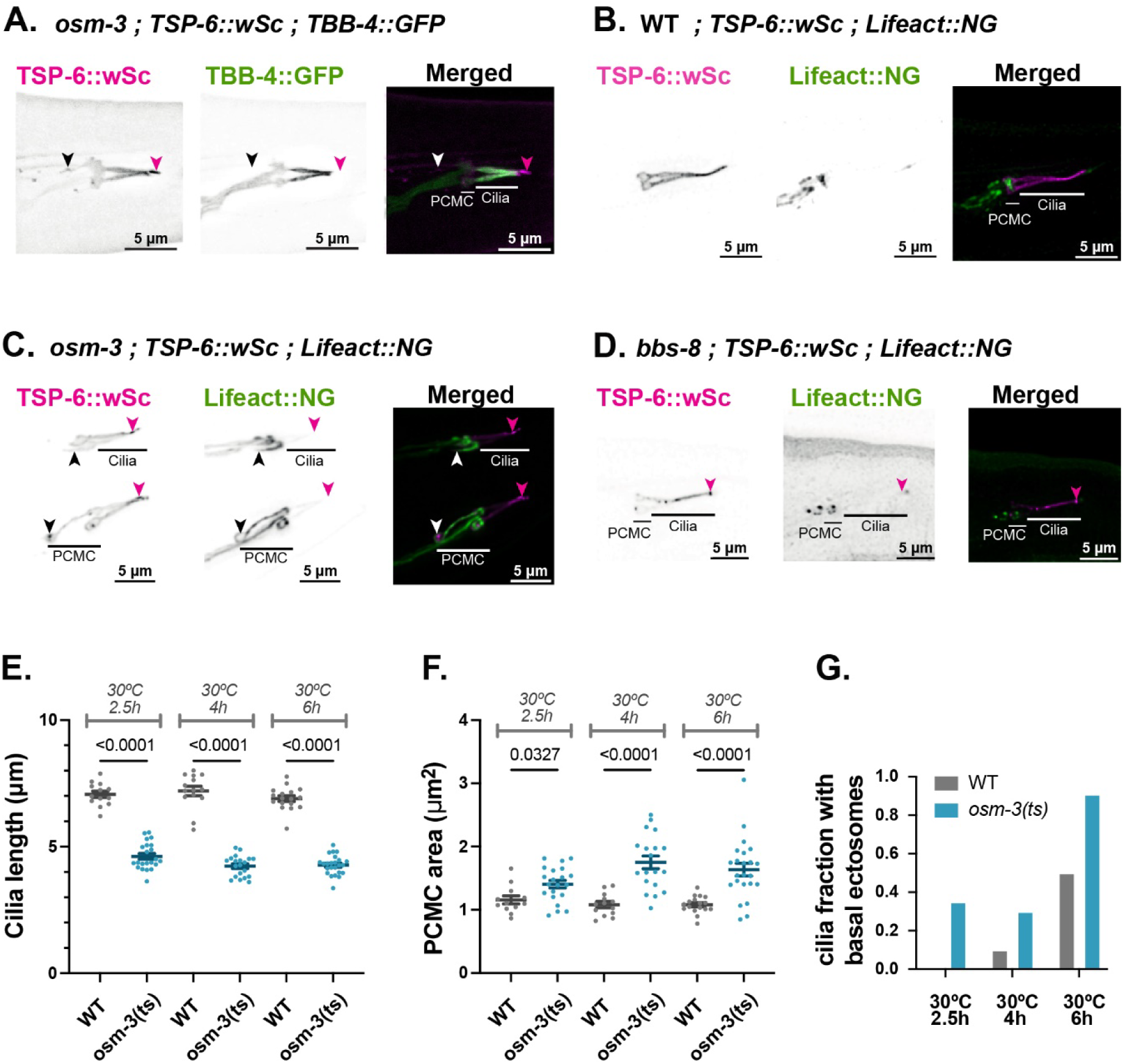
Cilia shortening is followed by PCMC enlargement, concomitant with basal ectosome accumulation in PHsh. (A) In *osm-3*, TBB-4::GFP is not observed within TSP-6::wSc positive PCMC branches (black arrowheads). TSP-6::wSc positive apical ectosomes (magenta arrowhead) budding from the cilia tip do are not enriched in TBB-4::GFP. (B) Lifeact::NG labeled F-actin is observed in the PCMCs and distal dendrite of PHA and PHB in N2. (C) Lifeact::NG is observed in the distal dendrite, PCMCs and the PCMC of *osm-3*, but not in apical ectosomes. (D) Lifeact::NG is observed in the distal dendrite and PCMCs of *bbs-8*, but not in apical ectosomes. (E) Shortening of the cilia proper is observed in *osm-3(ts)* after 2,5h of exposure to 30°C. (N comprised between 13 and 24). (F) The PCMC size of *osm-3(ts)* significantly increases after 4 h exposure to 30°C. (N comprised between 13 and 23). Brown– Forsythe ANOVA was performed followed by Dunnett’s T3 post-hoc test to correct for multiple comparisons. (G) After exposure to 30°C for 4h and 6h, basal ectosomes are released and detected in PHsh cytoplasm surrounding the cilia of *osm-3(ts)* (N comprised between 10 and 17). Brown–Forsythe ANOVA was performed followed by Dunnett’s T3 post-hoc test to correct for multiple comparisons.

Previously published ultrastructure of *osm-3* cilia suggested that microtubules were ectopically present within PCMC branches^66^. To assess microtubule localization, we imaged the axoneme-enriched beta-tubulin marker TBB-4::GFP. In *osm-3*, TBB-4::GFP was absent from PCMC branches indicating a lack of microtubules in these structures (Figure 7A). As the PCMC branches were not supported by a microtubule backbone, we hypothesized that F-actin might participate in PCMC extensions. The distribution of F-actin was examined using the F-actin marker Lifeact::NG. This marker was predominantly observed within the PCMC and distal dendrites of wild-type animals (Figure 7B). In the *osm-3* mutant, Lifeact::NG was enriched at the PCMC and its branches but rarely enter basal ectosomes, although these are derived from the PCMC and its branches (Figure 7C, Figure S4A). As ciliary F-actin was suggested to contribute to apical ectocytosis in BBSome mutants, we checked for the presence of Lifeact::NG in the cilia of *bbs-8* mutant animals^42,68^. Lifeact::NG was observed in the PCMC of *bbs-8* animals but neither in distal ciliary segment nor in apically derived ectosomes, suggesting F-actin does not invade the *bbs-8* cilia (Figure 7D).

Because ciliogenesis defects are observed in *osm-3*, *osm-6, che-11,* and *ift-74* mutants, we asked whether PCMC expansion and basal ectosome release observed in PHA and PHB mutant cilia are caused by developmental defects. To assess this, we induced acute ciliary shortening in adults using *osm-3(oy156)*: a temperature-sensitive *(ts)* allele of *osm-3*^69^. When raised at 15°C and imaged at 20°C, the cilia length and PCMC area of *osm-3(ts)* were indistinguishable from wild-type controls^69^. Transferring adult animals to a restrictive temperature (30°C) for 2,5 hours resulted in cilia shortening, and this phenotype persisted for at least 6 hours as the animals were kept at the restrictive temperature (Figure 7E). We sought to understand whether cilia shortening correlated with PCMC expansion and basal ectosome release. After a 4 hours temperature shift, significant PCMC enlargement was observed and basal ectosomes appeared in PHsh glia, in the vicinity of the PCMC (Figure 7F-G). These results demonstrate that PCMC expansion and basal ectosome release are dynamic events that can occur in response to acute changes in IFT and/or cilium shortening and are not a result of a developmental ciliogenesis defects.

## DISCUSSION

In this study, we established TSP-6::wSc as a ciliary membrane marker to investigate the process of ciliary ectosome biogenesis (ectocytosis). We have shown that TSP-6::wSc vesicles are actively transported along dendrites towards the cilium base. Once loaded at the PCMC, TSP-6::wSc moves along the ciliary membrane by lateral diffusion, independently of IFT. Some members of the tetraspanin protein family have been shown to form oligomeric complexes with themselves, other membrane proteins, and specific lipids^70,71^, but we did not observe the tetraspanin TSP-6::wSc to form such oligomeric complexes within the cilium. In the proximal segment of the wild-type cilium, the diffusion coefficients of the 50 kDa lipid-anchored aARL-13::NG and 53 kDa four-pass transmembrane protein TSP-6::wSc are comparable, suggesting that both markers diffuse freely within the membrane with minimal -if any-interactions with other ciliary components. However, in the distal segment, the diffusion coefficient of TSP-6::wSc is decreased, while that of aARL-13::NG remains high, suggesting differences in their molecular motion. Spontaneous partition of transmembrane proteins can occur owing to differences in their local molecular motion^72,73^. Accordingly, our simulations suggest that enrichment of TSP-6::wSc in distal cilia can be explained by its reduced diffusivity in the distal cilia. Because TSP-6::wSc does not form oligomeric complexes, its reduced diffusion coefficient in the distal cilium is more likely attributable to factors slowing down TSP-6::wSc -but not aARL-13-NG-such as molecular crowding, lipid composition, or the pronounced membrane curvature. Currently, neither the precise density of proteins nor the distribution of lipids along the ciliary membrane have been characterized. However, we know that membrane curvature is the highest in the ∼100 nm diameter distal segment of the cilium, where TSP-6::wSc is the most enriched (Figure S4B)^74^. Furthermore, conical transmembrane proteins tend to exhibit reduced diffusion under conditions of high membrane curvature^75^. Because of its conical structure, the orthologous tetraspanin CD9 has been shown before to passively sort itself into highly curved regions of the membrane, contributing to membrane remodeling and EV biogenesis^76,77^. Therefore, the retention of TSP-6::wSc in the distal cilium, membrane buds and ectosomes may be influenced by its partitioning towards highly curved membrane regions.

Mutations that alter the distribution of ciliary transmembrane proteins along cilia – such as in the IFT mutants-are also expected to modify the inclusion of these proteins into apical, proximal or basal ectosomes. TSP-6::wSc has unique properties: it distributes along the PCMC and the cilium by lateral diffusion within the ciliary membrane independently of IFT or TZ and enters cEVs. These characteristics make it a reliable marker for measuring ectocytosis rates, regardless of the mutant background or the budding site along the cilium. In control conditions, we observed that ectocytosis rates appear low for PHA and PHB compared to male ray neurons or CEMs and for the sex-shared IL2, which can produce nearly 400 ectosomes per hour in control conditions^27,28,30^ Increased rate of ectosome production has previously been observed from the male cilia of *inpp-1, bbs-8* or *nphp-4* mutants^28,30,78^. Similarly, the ectosome release rate was increased in PHA and PHB in *tub-1, bbs-8, arl-13* mutants known to participate in ciliary transmembrane protein trafficking and in *pdl-1*, *inpp-1*, *cil-1* mutants known to control ciliary PI(4,5)P_2_ distribution^9,12,14,37,41,79^. In addition, we observed that ectocytosis takes place in different parts of the cilium, depending on the nature of the mutation. Remarkably, the location of ectocytosis appears to correlate with the ciliary compartment in which transmembrane proteins accumulate in these mutants. In the absence of *tub-1* (TULP3 in mammals), which mediates the ciliary entry of transmembrane proteins, we observed enhanced basal ectocytosis from the PCMCs where accumulation of GPCRs and cGMP-gated channel occurs^9,12^. The BBSome promotes ciliary retrieval and degradation of specific cargo proteins^13,14,40,80^, and increased apical ectocytosis of GPCR, polycystin or guanylate cyclases were previously observed in mutants where retrieval by the BBSome was compromised^24,42^. In absence of the BBSome subunit *bbs-8*, we also observed enhanced ectocytosis from the cilium proper, where accumulation of GPCRs, polycystins, and guanylate cyclases were previously observed^40,41,81^. GPCRs have been shown to accumulate in the cilium proper of a INPP5E mouse mutant, likely because the high ciliary PI(4,5)P_2_ concentration disturbs GPCR trafficking by TULP3/TUB-1^11,82,83^. Accordingly, we observed increased apical ectocytosis in mutants with increased ciliary PI(4,5)P_2_: *pdl-1, inpp-1, cil-1, mks-5*. Finaly, ARL13b is thought to facilitate the retrieval of specific transmembrane-protein cargoes by coupling them to the BBSome^17,18^. In *arl-13* mutants, we observed ectosome production from the proximal segment of the cilia where GPCRs, polycystins, and cGMP-gated channels have been shown to accumulate^9,17^. Altogether, these observations suggest that buildup of ciliary transmembrane proteins into microdomains of the ciliary membrane induces or supports local membrane budding and ectosome formation, facilitating the removal of these transmembrane proteins from the cilia. Ciliary transmembrane proteins include receptors, but also adhesion proteins, enzymes involved in post-translational modifications, lipid-modifying enzymes, and others^31^. In contrast to transmembrane proteins, the location of soluble protein buildup in the ciliary cytoplasm appears less determinant: we observed that *ift-74* and *che-11* causes distal cilia buildup of IFT-A complexes but associates with basal ectocytosis.

We observed that acute exposure to sensory cues induced cilia remodeling together with ectocytosis. Within one minute after cue delivery, IFT is interrupted, the distal axoneme collapses, fission of the ciliary membrane occurs as proteins present in the distal cilium appeared to be captured within apical ectosomes and are subsequently ejected through the phasmid channels into the environment. Interestingly, basal ectosome release was observed ∼20 min following cilium retraction, in animals still lacking the distal segment of the cilia. Among the ciliary pathways potentially involved in short and long-term cilia remodeling and ectocytosis, our results implicate EGL-4/PKG activity and the spatial distribution of the phosphoinositide PI(4,5)P_2_ lipid. Increased apical ectocytosis in *pdl-1, inpp-1* and *cil-1* suggests that increased ciliary PI(4,5)P_2_ levels promote ectocytosis from the cilia. Reduced basal ectocytosis in the absence of the PDL-1 shuttle suggests that reduced PI(4,5)P_2_ levels in the PCMC prevent basal ectocytosis.

In the ciliary mutants that we studied here, we observed a striking anticorrelation between ciliary length and PCMC size (Figure S4C). A large PCMC correlated with high rate of basal ectosome production, while elongated cilia correlated with apical ectosome production (Figure S4D). Therefore, we envision a mechanism that couples ciliary architecture to ectocytosis location. A transient expansion of the distal cilia prior to ectocytosis may explain why longer cilia are associated with apical ectocytosis, as we observed in timelapse imaging of *mks-6* mutants. In contrast, mutants displaying shorter cilia correlated with larger PCMC and increased basal ectosome production (Figures 6F, Figure S4C). In *osm-3(ts),* the cilia retracted first prior to PCMC expansion and basal ectocytosis suggesting that cilium shortening may promote PCMC expansion and basal ectocytosis. Potential mechanisms coupling cilium shortening with PCMC expansion and basal ectocytosis include the disruption of membrane import/export balance, the redistribution of IFT and cargoes from the distal cilia to the proximal cilia and PCMC, and the disruption of ciliary signaling. Although membrane import/export to/from the PCMC influences cilium architecture, the mechanisms remain poorly explored^60,63,84,85^. We observed increased apical ectocytosis in *cav-1* mutants or *dyn-1(K46A)* transgenic animals that reduce endocytosis rate, suggesting membrane import/export balance to/from the PCMC can influence distal cilium mechanisms.

Among all transmembrane proteins whose buildup may initiate ectocytosis, our results do not support a direct role for TSP-6. Indeed, IFT interruption should not directly affect the distribution of TSP-6, which occurs via normal diffusion within the ciliary membrane. However, transmembrane proteins buildup occurs in the ciliary membrane during transient IFT interruption or in *tub-1*, *bbs-8*, or *arl-13* mutants. We suggest that protein crowding in microdomains of the ciliary membrane may cause secondary retention of TSP-6 at buildup sites, because of TSP-6’s reduced molecular motion/ diffusion rate. Accordingly, TSP-6-enriched membrane buds are observed prior to ectocytosis. Similarly to CD9, the conical shape of TSP-6 may promote positive membrane curvature ^76,77^. A positive feedback loop could then drive increased membrane curvature by spontaneous partitioning of conical-shaped TSP-6 to curved regions of the membrane, ultimately leading to TSP-6-enriched membrane buds. In addition, TSP-6 partitioning may favor the capture of other transmembrane proteins into membrane buds because of local protein crowding. In this line, in the *bbs-8* mutant cilia, the TRPV channel OCR-2::eGFP enters membrane buds enriched with TSP-6::wSc along the cilia and also enters into apical ectosomes. Interestingly, this model would spontaneously sort the crowded proteins to ectosomes.

In conclusion, using a diffusive ciliary membrane marker for ectocytosis, we show that sensory cues interrupting IFT also induce acute apical ectocytosis, clearing out material from the distal segment of the cilium. Mutants causing distal, proximal or periciliary buildup of transmembrane proteins buildup increase distal, proximal or periciliary ectosome biogenesis, respectively. Therefore, local transmembrane receptor accumulation and ectocytosis from these sites appear to be coupled. Ciliary transmembrane receptors are dynamically gated by their activity^13,14,42,80,86,87^. In that context, the same mechanism may selectively remove the buildup of receptors by ectosomes during cilia signaling or when ciliary trafficking is disrupted.

## ACKNOWLEDGEMENTS

We thank Dr. Aniruddha Mitra and Dr. Noémie Danné (Vrije Universiteit Amsterdam) for help with the data analysis. We thank J.M.Vanderwinden and LiMiF for the help with confocal microscopy. We thank P. Sengupta lab for the *osm-3(ts)* strain. We acknowledge financial support from the European Research Council under the European Union’s Horizon 2020 research and innovation programme (grant agreement no. 788363; “HITSCIL”; to EJGP), the Dutch Research Council (NWO; Project no. OCENW.M20.063 to GH and EJGP), the Belgian Fonds de la Recherche Scientifique-FNRS (PDR 40020876 to PL, FRIA 1.E.098.24F to AN, 1.E.065.22F to TL). Some strains were provided by the CGC, which is funded by NIH Office of Research Infrastructure Programs (P40 OD010440).

## AUTHOR CONTRIBUTIONS

Conceptualization, T.L., G.H., A.N., C.B., A.R., P.L., E.P. Methodology, T.L., G.H., A.N., C.B., A.R. Analysis and figures T.L., G.H., A.N., C.B. Investigation, T.L., G.H., A.N., C.B. and A.R. Writing – original draft, P.L. Writing – review & editing, T.L., G.H., A.N., C.B., A.R., P.L., E.P. Funding acquisition, P.L, E.P.

## MATERIALS AND METHODS

### C. elegans maintenance

For all experiments, *C. elegans* were grown on NGM plates, synchronized using an egg-laying window, and reared at 20 °C in the presence of the OP50 bacterial strain until the first day of adulthood (D1). The wild-type control strain corresponds to the TSP-6::wScarlet knock-in made in the N2 genetic background. All the other strains are described in Table 1.

**Table 1:** Strains list.

| **Strain** | **Description** | **Source** |
| --- | --- | --- |
| PHX4122 | tsp-6(syb4122[Ptsp-6::tsp-6::wrmScarlet]) X. | <sup>24</sup> |
| OQ366 | tsp-6(syb4122[Ptsp-6::tsp-6::wrmScarlet]) X.; ulbEx78 [pF16F9.3::CFP] | <sup>24</sup> |
| OQ868 | Ex[pGpa-11::StG::RAB-8] ;OQ366 | This study |
| OQ872 | Ex[pOsm-10::AMAN-2::StG]; OQ366 | This study |
| OQ871 | Ex[pGpa-11::StG::RAB-5]; OQ366 | This study |
| OQ870 | Ex[pGpa-11::StG::RAB-11]; OQ366 | This study |
| OQ869 | Ex[pGpa-11::StG::RAB-7]; OQ366 | This study |
| OQ873 | Ex[pOsm-3::LMP-1::EGFP]; OQ366 | This study |
| DR1 | unc-101(m1) I. | CGC |
| OQ801 | unc-101(m1) I.; OQ366 | This study |
| OQ888 | Ex[pOsm-3::arl-13::NG] ; OQ366 | This study |
| OQ885 | Ex[pOsm-3::lifeact::NG] ; OQ366 | This study |
| OQ886 | bbs-8(nx77) V.; Ex[pOsm-3::lifeact::NG] ; OQ366 | This study |
| OQ887 | osm-3(p802) IV.; Ex[pOsm-3::lifeact::NG] ; OQ366 | This study |
| EJP501 | vuaSi001 [Pocr-2::ocr-2::eGFP] IV | <sup>96</sup> |
| EJP76 | vuaSi15 [pBP36; Posm-6::osm-6::eGFP; cb-unc-119(+)] I; unc-119(ed3) III; osm-6(p811) V | <sup>90</sup> |
| EJP212 | vuaSi26 [pJM6; Pxbx-1::xbx-1::EGFP; cb-unc119(+)] I; vuaSi2[pBP22;Posm-3::osm3::mCherry; cb-unc-119(+)] II; osm-3(p802) IV; xbx-1(ok279) V | <sup>97</sup> |
| DG2179 | tub-1(nr2044) II. | CGC |
| OQ533 | tub-1(nr2044) II.; OQ366 | This study |
| OQ368 | bbs-8(nx77) V.; OQ366 | <sup>24</sup> |
| OQ813 | tbb-4-GFP(KI) X.; PHX4122 | This study |
| OQ814 | osm-3(p802) IV.; tbb-4-GFP(KI)X.; PHX4122 | This study |
| OQ815 | bbs-8(nx77) V.; tbb-4-GFP(KI) X.; PHX4122 | This study |
| MT1074 | egl-4(n479) IV. | CGC |
| OQ784 | egl-4(n479) IV.; OQ366 | This study |
| DA521 | egl-4(ad450) IV. | CGC |
| OQ840 | egl-4(ad450) IV.; OQ366 | This study |
| OQ369 | osm-3(p802) IV.; OQ366 | <sup>24</sup> |
| PR811 | osm-6(p811) V. | CGC |
| OQ793 | osm-6(p811) V.; OQ366 | This study |
| CB3330 | che-11(e1810) V. | CGC |
| OQ790 | che-11(e1810) V.; OQ366 | This study |
| FX2394 | ift-74(tm2394) II. | CGC |
| OQ786 | ift-74(tm2394) II.; OQ366 | This study |
| RB2574 | mks-5(ok3582) II. | CGC |
| OQ785 | mks-5(ok3582) II.; OQ366 | This study |
| VC1466 | mks-6(gk674) I. | CGC |
| OQ498 | mks-6(gk674) I.; OQ366 | This study |
| PT709 | <u>nphp-4(tm925)</u> <u>him-5(e1490)</u> V. | CGC |
| OQ837 | nphp-4(tm925) V.; OQ366 | This study |
| VC282 | pdl-1(gk157) II. | CGC |
| OQ824 | pdl-1(gk157) II.; OQ366 | This study |
| CX4544 | ocr-2(ak47) IV. | CGC |
| OQ844 | ocr-2(ak47) IV.; OQ366 | This study |
| VC1073 | rabs-5(ok1513) IV. | CGC |
| OQ495 | rabs-5(ok1513) IV.; OQ366 | This study |
| RB1679 | cav-1(ok2089) IV. | CGC |
| OQ792 | cav-1(ok2089) IV.; OQ366 | This study |
| OQ843 | Ex[posm-10::dyn-1(K46E)::SL2GFP] ; OQ366 | This study |
| VC2109 | Y53F4B.4&Y53F4B.42(gk1068) II; T25B9.10(gk3262) IV;<br>K04C1.5(gk3263) X. | CGC |
| OQ417 | inpp-1(gk3262) IV.; OQ366 | This study |
|  | cil-1(tm3568) III. | NRBP |
| OQ835 | cil-1(tm3568) III.; OQ366 | This study |
| VC1104 | Y37E3.4&Y37E3.5(gk513) I. | CGC |
| OQ796 | arl-13(gk513) l.; OQ366 | This study |
| PY12081 | osm-3(oy156) | Gift from<br>Sengupta P. |
| OQ493 | osm-3(oy156) ; OQ366 | This study |

### Analysis of the cilia morphology and basal ectosome release quantifications

Between 10 to 20 synchronized D1 adult animals were placed on a 4% agarose pad with 10 µl of 10 mM levamisole in M9 buffer for immobilization. Imaging was performed within one hour of anesthetizia using a LSM780NLO confocal system on an inverted microscope (Carl Zeiss, Oberkochen, Germany). Two objective lenses were used: either LD C-Apochromat 40x/1.1 W Korr M27 or alpha-plan-Apochromat 63x / 1.46 Oil Korr M27 (both from Carl Zeiss). The TSP-6::wScarlet signal was imaged with 543 nm laser excitation and detected between 570–695 nm wavelengths. The PHsh::CFP signal was imaged with 458 nm excitation and detected between 463-558 nm, Single channel images were acquired using GaAsP detector. For dual channel imaging, wScarlet was collected with the GaAsP detector and the CFP signal was collected simultaneously using the PMT detector. Confocal images were processed using FIJI^88^.

To characterize cilia morphology, entire cilia were imaged using a 63x / 1.46 Oil Korr M27 objective 368 x 368 pixels frame, with a pixel size of 0.09 μm × 0.09 μm. The pinhole was set to 0,93 Airy Units, the Z-step interval to 0.41 μm and pixel dwell time to 1.11 μs. Z-stack acquisitions were processed into 2D images using maximum intensity projection to generate flattened representations of the 3D volumes. To determine the PCMC size, the area was measured by manually outlining the region with the polygon selection tool. Cilia length was assessed by drawing a line from the base of the PCMC to the tip of the cilium using the segmented line tool.

To characterize basal ectosome release, we acquired the whole PHsh Soma using a 464 x 464 pixels frame with a pixel size of 0.10 μm × 0.10 μm. The pinhole size was set to 0,74 Airy Units for wScarlet detection and 0,92 Airy Units for CFP detection. Z-stacks were acquired with a step size of 0.52 μm and a pixel dwell time of 1.41 μs. Z-stack acquisitions were processed into 2D images using maximum intensity projections in FIJI, generating flattened representations of the 3D volumes. PHsh fluorescence intensity was used to quantify the production of basal ectosome over a period longer than 24h. A region of interest (ROI) was defined by manually outlining the PHsh cell body using the polygon tool. Measurements of area, integrated density, and mean gray value were recorded. To assess the TSP-6 signal within the ROI, the corrected total cell fluorescence (CTCF) was calculated using the formula: CTCF = Integrated Density – (Area of ROI × Mean Gray Value). To account for variations in PHsh size, the CTCF was normalized by dividing it by the ROI area.

All basal cilia ectosome measurements were obtained from three independents imaging sessions, each including WT controls and mutant samples. Prior to pooling, consistency across sessions for WT controls and mutants was assessed statistically to confirm comparability. Statistical analysis and graph design were performed in Prism GraphPad Prism (GraphPad Software, San Diego, CA, USA).

### Acute morphological changes induced in by heat in *osm-3(oy156)*

For all experiments that include *osm-3(oy156)*, all worms were synchronized by an egg-laying window and reared at 15°C up to D1. Prior to imaging, each group of D1 was incubated at 30°C for 2 h, 3.5 h or 5.5 h before mounting, and an additional 30 minutes at 30°C after mounting. ∼10 animals were placed in 10 µl of 10mM levamisole dissolved in M9 buffer and mounted on a 2% agarose pad. All images were acquired on the Inverted Zeiss LSM 780 confocal system using 63x / 1.46 Oil Korr M27 objective. Excitation (543 nm laser) and detection (570–695 nm) wavelengths were adjusted to capture the TSP-6::wScarlet signal acquired using GaAsP detector. The settings for these images were as follows: frame size was set to 512 x 512 pixels with a pixel size of 0.05 μm × 0.05 μm, pinhole was set to 2.5 Airy Units, Z-step to 0.4 μm/step, pixel dwell to 0.79 μs, and averaging to 2. Images from 2 separate imaging sessions were analyzed using FIJI. Statistical analysis and graph design were performed in Prism GraphPad.

### Apical ectosome quantification assay

Coverslips were coated by placing 150 µl of the 0.01% poly-L-lysine solution (#A-005-M, Merck) per coverslip and letting coverslips dry overnight in a hood. ∼10 synchronized D1 adults were placed in a 10 µl droplet of 10 mM levamisole dissolved in M9 buffer on the poly-L-lysine-coated coverslips. After applying vaseline on the corners of the coverslips, microscopic slides with a 5% agarose pad were placed over the animals. Images were acquired one hour after anesthesia. All images were acquired on the Inverted Zeiss LSM 780 confocal system using 63x / 1.46 Oil Korr M27 objective. Excitation (543 nm laser) and detection (570–695 nm) wavelengths were adjusted to capture the TSP-6::wScarlet signal acquired using GaAsP detector. PHA/PHB neurons cilia were placed in the center of imaging ROI and Z-stack were acquired as follows: frame size was set to 512 x 512 pixels with a pixel size of 0.26 μm × 0.26 μm, pinhole was set to 12 Airy Units, Z-step to 0.4 μm/step size, pixel dwell to 1.27 μs, and averaging to 1. Acquisition was performed in 3 separate imaging sessions on different days, each including all analyzed mutants and the wild-type control strain (N2), to demonstrate consistent phenotypes across sessions.

Z-stack acquisitions were converted into a 2D image using maximum intensity projections in FIJI to obtain a flattened image representative of the 3D volume. Released apical ectosomes were quantified by counting them manually in the acquired ROI. If vesicles formed a cluster making them inadequate to count, the image was excluded from analysis. Results from the 3 separate sessions were pulled together. Statistical analysis and graph design were performed in Prism GraphPad.

### Time-lapse imaging of ectosome release in *osm-3, mks-6* and *arl-13* mutants

Around 20 synchronized D1 adult animals were placed on a 2% agarose pad with 10 µl of 10 mM levamisole in M9 buffer. Time-lapse images were acquired within 30–60 minutes after anaesthesia.

The acquisition settings for *osm-3* were as follows on an Inverted Zeiss LSM 780 confocal system using 63x / 1.46 Oil Korr M27 objective. Excitation (543 nm laser) and detection (570–695 nm) wavelengths were adjusted to capture the TSP-6::wScarlet signal acquired using GaAsP detector. Frame size was set to 512 x 512 pixels with a pixel size of 0.03 μm × 0.03 μm, pinhole size was set to 1 Airy Unit, Z-step to 0.41 μm/step size, pixel dwell to 1.27 μs, and averaging to 4. Each Z-stack made of 3 Z-step was taken every 5 s. Before starting the time-lapse, 1 dual-channel Z-stack was acquired to align with single-channel time-lapse. In addition to TSP-6 acquisition channel, the excitation (458 nm laser) and detection (463–558 nm) wavelengths of the second channel were adjusted to capture the PHsh::EGFP signal.

The acquisition settings for *mks-6* images were as follows on an Inverted Zeiss LSM 780 confocal system using 63x / 1.46 Oil Korr M27 objective. Excitation (543 nm laser) and detection (570–695 nm) wavelengths were adjusted to capture the TSP-6::wScarlet signal acquired using GaAsP detector. Frame size was set to 300 x 300 pixels with a pixel size of 0.05 μm × 0.05 μm, pinhole size was set to 1 Airy Unit, Z-step optical sections 0.2 μm/step size, pixel dwell was set to 2.7 μs, and averaging was set to 1. Each Z-stack made of 5 Z-step was taken every 6,5 s.

The acquisition settings for *arl-13* images were as follows on an Inverted Zeiss LSM900 Airyscan confocal system using 63x / 1.40 Oil objective. Excitation (543 nm laser) and detection (570–695 nm) wavelengths were adjusted to capture the TSP-6::wScarlet signal acquired using Airyscan super resolution detector. Frame size was set to 322 x 322 pixels with a pixel size of 0.049 μm × 0.049 μm, pinhole was set to 5 Airy Units, pixel dwell to 3.06 μs, and averaging to 1.

All the time-lapse images were processed using FIJI. For super resolution images, Airyscan processing was applied beforehand. Z-stack acquisitions were converted into a 2D image using maximum intensity projections to obtain a flattened image representative of the 3D volume.

### Colocalization analysis

Around 20 synchronized D1 adult animals were placed on a 2% agarose pad with 10 µl of 10 mM levamisole in M9 buffer. Images of the neuronal soma and cilia were acquired in the following 30-60 min after animals were anesthetized. To assess the colocalization of TSP-6::wSc with different markers (AMAN-2::StG, StG::RAB-8, StG::RAB-5, StG::RAB-11, StG::RAB-7, LMP-1::GFP, OCR-2::GFP), images were acquired on a Zeiss LSM900 Airyscan confocal system, 63x/1.40 oil objective. Excitation (488 and 543 nm laser) and detection (490-560 and 570–695 nm) wavelengths were adjusted to capture TSP-6::wScarlet and StG/GFP signals, respectively using Airyscan super resolution detector. The acquisition settings were as follows: frame size was set to 795 x 795 pixels with a pixel size of 0.043 μm × 0.043 μm, pinhole was set to 5 Airy Units, pixel dwell to 2.66 μs, and averaging to 2. StG::RAB-8, StG::RAB-5, StG::RAB-11, StG::RAB-7 images were acquired in the ASH/ASI neuron, whereas AMAN-2::StG and OCR-2::GFP images were acquired in the PHA/PHB neurons.

After Airyscan processing, all images were analyzed using FIJI. Supplementary videos present the whole acquired z-stack in which each frame presents one z-step. Colocalizing spots are highlighted by arrows. Images of AMAN-2::StG and OCR-2::GFP show 1 selected Z-step as a representation of the whole Z-stack.

### Single-molecule microscopy with SWIM

Single-molecule imaging was performed with a custom-built epi-illuminated fluorescence microscope^89,90^ and using small-window illumination microscopy (SWIM)^91^. This setup uses an inverted microscope body (Nikon Ti E) equipped with a 100x oil immersion objective (Nikon, CFI Apo TIRF 100x, N.A.:1.49). The excitation source in this system included a 491 nm (Cobolt Calypso, 50 mW) and 561 nm (Cobolt Jive, 50 mW) DPSS lasers. The wavelength and laser power were controlled using an acousto-optic tunable filter (AOTF, AA Optoelectronics). To narrow the beam size for SWIM, an iris diaphragm (Thorlabs, SM1D12, ø 0.8-12 mm) was positioned in the focal plane of the epi lens. The size of the aperture was adjusted manually to switch between the small illumination beam for SWIM and the full beam for normal imaging. Inside the microscope body, the fluorescence was separated from the excitation light by a dichroic mirror (ZT 405/488/561 rpc; Chroma) and appropriate emission filter for the wScarlet channel (525/45; Brightline HC, Semrock) or the NeonGreen Channel (525/45; Brightline HC, Semrock). Images were then recorded on an EMCCD camera (Andor, iXon 897). The whole system was controlled using MicroManager software (v1.4) (Edelstein, 2010).

To achieve single-molecule imaging, we illuminated a 10-15 µm size window centered around a pair of cilia. After prolonged illumination with high laser power, the single-molecule regime is reached due to photobleaching of most fluorescently labeled molecules, after which the laser power was decreased to sustain this regime before all molecules were photobleached. All single-molecule images were obtained with a frame rate of 20 Hz. Since the influx of new TSP-6::wSc molecules in wild-type and mutant strains into the cilium is very low, it was only possible to remain in the single-molecule regime for about 5 min. In the case of aARL-13::NG it was possible to do single-molecule imaging for more than 10 min.

### Line-scan confocal microscopy

For the two color fluorescent image in Figure 1A and the kymograph in Figure 1E in which the entire PHA or PHB neuron was imaged for long periods of time, a line-scan confocal microscope was used (Confocal.nl, NL5+). The confocal unit is attached to an inverted microscope body (Nikon Ti E) equipped with a 60x water immersion objective (Nikon, CFI Plan Apo IR, NA 1.27). To enable fast Z-stacks, the microscope was also equipped with a piezo stage (MCL, nanodrive). The excitation light is provided by a four-color laser engine (Toptica, iChrome CLE-50) and fiber-coupled to the confocal unit. Images were recorded by a sCMOS camera (Hamamatsu, Orca Flash v2.0).

### Acute chemical exposure with microfluidic worm chip

We used a microfluidic chip^92^ to accurately expose the tail of the worm to chemical repellents, such as 0.1% sodium dodecyl sulfate (SDS) or 10 mM copper sulfate (CuSO4). This microfluidic design has four input channels, including a buffer channel, the repellent and two channels filled with a fluorescent dye (fluorescein, Merck). The latter was used to visualize the flows under the microscope and calibrate them. In these experiments M13 was used as a buffer and to dilute the repellent and fluorescein. The input channels of the microfluidic chip were connected to 3 ml syringes pressurized using a flow control system (Fluigent, MFCS-EZ) and controlled with the included software (Fluigent All-in-one 2019). Before to insert them in the microfluidic chip, the worms were anesthetized for 15 minutes in a droplet of 5 mM Levamisole. A single worm was aspirated into a Tygon tube via a stainless-steel pin (0.025” OD, 0.013” ID; New England Small Tube Corporation) connected to a 3 mL syringe containing M13 buffer. The worm was then injected tail-first into the worm trap through the worm channel inlet by manual syringe pressure. A detailed description of this setup and protocols, calibration and usage can be found in^38,93^.

The microfluidic system was installed on a near identical microscope setup as used in the single-molecule experiments described above. Therefore, we were able to do long term time-lapse imaging before and after exposure to chemical repellents as well as the SWIM protocol to track TSP-6 vesicles and extract their motion characteristics in the dendrite or their interaction with the PCMC.

### Tracking and Analysis single-molecule TSP-6

Single-molecule tracking was performed using FIESTA (version 1.6.0), a MATLAB-based program ^94^. Each track, representing a single-molecule event, contained time, x and y coordinates, and distance traveled for each frame. Post-processing was done using an overlay of the connected tracks on a kymograph. Only tracks corresponding to single-molecule events located in the PCMC, TZ, or cilium proper, and lasting at least 12 frames, were included. Tracks resulting from closely overlapping single-molecule events (and thus indistinguishable) were excluded from analysis.

After tracking, the x and y coordinates of the tracks were converted to ciliary coordinates, which allowed pooling data from cilia across multiple worms. Using a custom Matlab script, a segmented line was drawn along the length of a cilium displayed as the time-averaged Z-stack of that single-molecule time-lapse movie. The conversion to ciliary coordinates was done by interpolating a cubic spline on this segmented line. Then, we chose the reference point at the intersection of the PCMC and the transition zone, as is forms an easily recognizable ‘bone-shaped’ structure consistent across samples that marks the start of the cilium. The conversion to ciliary coordinates resulted in all single-molecule localizations to be transformed from x and y to distance from the reference point along the spline and distance perpendicular to the spline^5^.

### Diffusion coefficient calculation

The local diffusion coefficients were determined by calculating the mean squared displacement (MSD) for time intervals of Δt, 2Δt, and 3Δt within a sliding window of 9 frames^95^. A linear fit was performed on the MSD versus time relationship, and the slope of the fit was taken as the local diffusion coefficient. This process was repeated for each position along the track. Finally, the diffusion coefficients calculated from the overlapping windows were averaged to provide a single diffusion coefficient for each point in the trajectory. This analysis was performed separately for displacements both parallel and perpendicular to the fitted spline. All calculations were performed using a custom-written MATLAB script.

### Point-to-point vesicle velocity

We used SWIM to image vesicles travelling in the dendrite as done before^91^. A circular beam with a diameter of about 15 µm was positioned right next to a pair of cilia, to avoid bleaching the fluorescent particles in the cilia that could become future retrograde vesicles. Tracking and spline fitting the trajectories of TSP-6::wSc vesicles in the dendrite was done using the same analysis pipeline as for the single-molecule data in the cilium. The point-to-point velocity was calculated by fitting a linear regression on the parallel distance traveled against the corresponding time points using a sliding window of 9 frames and taking the slope. After filtering (explained below) we determined the mean and standard deviation of the distribution.

Since the transport of TSP-6:wSc vesicles in the dendrite was bidirectional and paused frequently, we needed to filter for the point-to-point velocities, in order to extract the velocities of episodes of purely directed transport to gain insight into the motor proteins driving this transport. This was done by extracting the anomalous exponent value (*α*) of the time lag (*τ*) from *MSD = 2 Γ τ* (where *Γ* is the generalized transport coefficient) for each datapoint by using the windowed Mean Square Displacement classifier (wMSDc) approach, as described by^95^. The window used to calculate the α was kept fixed at 15 data points. Thus, trajectories shorter than 15 data points were disregarded. On top of that, only trajectories that showed active transport were selected, pausing and diffusive trajectories were excluded. We used an *α*-value of 1.7 to filter the velocity data. This value was chosen arbitrarily to include as much of the data as possible while at the same time excluding episodes of diffusive motion.

### Vesicle frequency

The vesicle frequency was determined by analyzing kymographs and manually selecting anterograde and/ or retrograde vesicles. The frequency was simply determined by dividing the number of vesicles by the total time of the measurement and repeated for multiple worms and in different worm strains. The statistical analysis and plotting were done with Prism GraphPad.

### Vesicle Intensity

The fluorescence intensity of each individual vesicle was estimated using Fiji^88^ by manually placing a box, large enough for all vesicles passing through, to capture the vesicle’s emitted fluorescence light. The box was positioned at the edge of the illumination spot where the vesicle would first appear, minimizing reducing the intensity by photobleaching. Therefore, anterograde and retrograde directed vesicles would be on either side of the illuminated spot. The intensity of vesicles traversing the box was recorded, and background intensity, measured from a box of the same size placed outside the dendrite, was subtracted.

### Bootstrapping

In some of the statistical analysis we employed a bootstrapping approach to estimate the average value and error of the distribution. This involved the random selection of N measurements from the distribution, performed with replacement, followed by the computation of the median for each resampled group. This procedure was iterated 1000 times, resulting in a bootstrapping distribution of medians. Subsequently, we calculated the mean (μ) and standard deviation (σ) of this distribution, which were then utilized to estimate the parameter and its associated error. With this bootstrapping method all reported values and errors are presented in the format μ ± 3σ.

### Diffusion simulation

We simulated the diffusion in the cilium to investigate whether a lower diffusion coefficient in the distal segment could explain the ciliary distribution of TSP-6::wSc we are observing. This was done by solving the 1D diffusion equation numerically using an explicit finite difference scheme. The one-dimensional diffusion equation, given by:

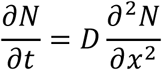

where N(x, t) represents the diffusing quantity, x is the spatial coordinate, t is time, and D is the diffusion coefficient, was solved numerically using a finite difference scheme. The spatial domain was discretized into M grid points with a uniform spacing of Δx = L/M, where L is the length of the cilium. The time domain was discretized into time steps of Δt.

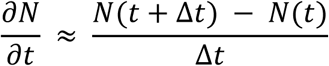

The central difference scheme was used to approximate the spatial derivative:

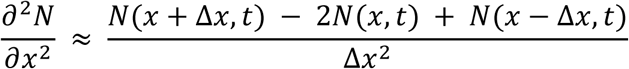

Combining those formulas gives us the number of molecules N depending on x and t.

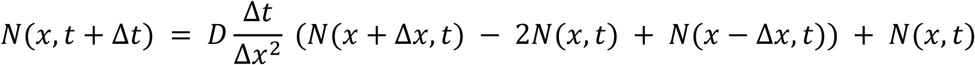

The cilium exhibits a decreasing diameter, and consequently, a decreasing perimeter towards its distal tip. We included this feature in the simulation by assuming that if one of the four contributions above involved a decrease in perimeter, a fraction of molecules proportional to the decrease would not diffuse but remain in the discretized grid point. Conversely, if it involved a perimeter increase, a proportional increase in the diffusive flux was applied, reflecting the augmented capacity for molecular transport in the expanded region.

The simulation was initialized with a fixed molecular concentration at the cilium base (x = 0) and a zero-flux reflective boundary condition at the distal tip. The spatial domain spanned 8 µm with a discretization of Δx = 0.1 µm, and the temporal discretization was Δt = 0.01 s. The diffusion coefficient was set to 0.2 µm²/s for simulations with a constant coefficient and varied linearly between 0.2 µm²/s and 0.1 µm²/s along the cilium’s length for simulations incorporating a spatially dependent diffusion coefficient.

We checked the stability of the method with the following condition:

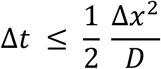

### Molecular biology

Expression vectors were generated by gateway LR reactions as described in Table1; let-858 3’UTR is present in the backbone of the destination vector, when not used in pentry pos3.

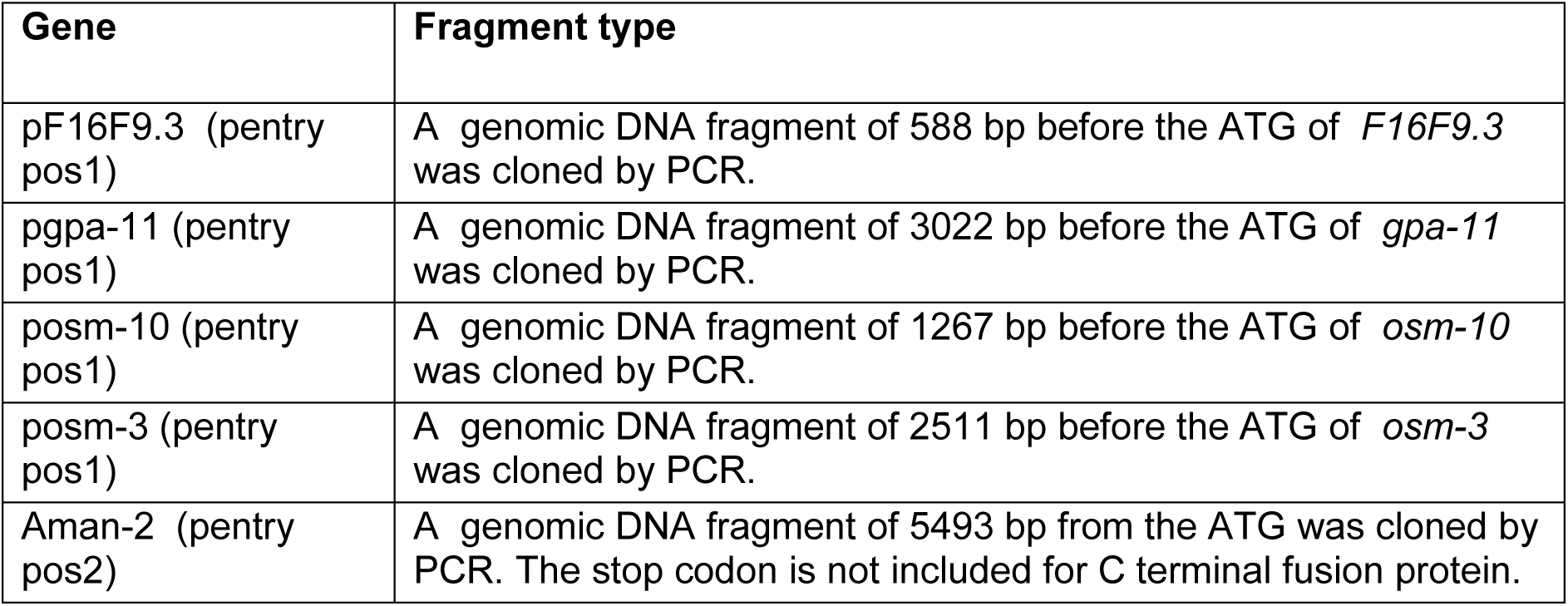
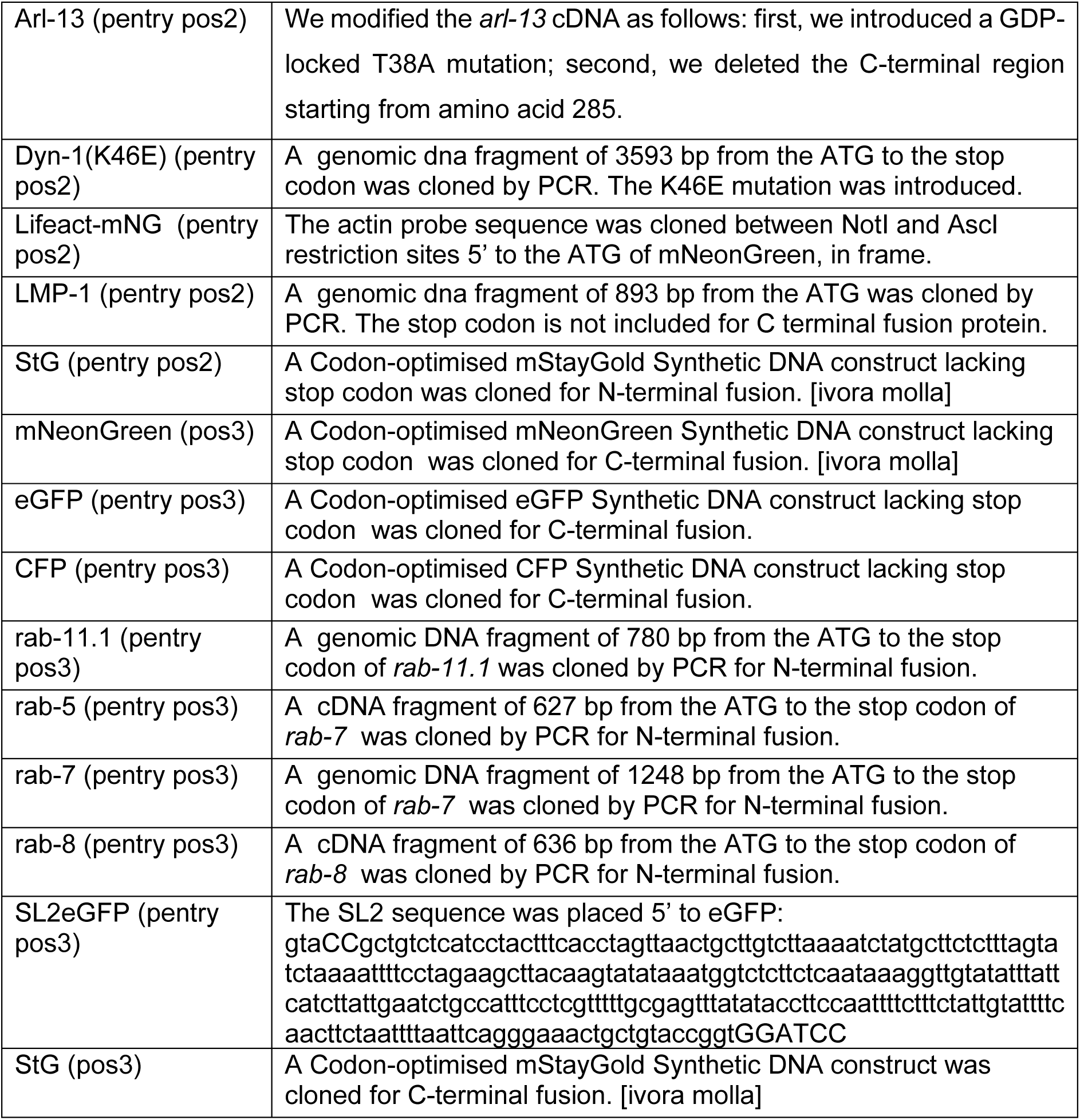

## SUPPLEMENTARY FIGURES

**Supplementary Figure 1:**
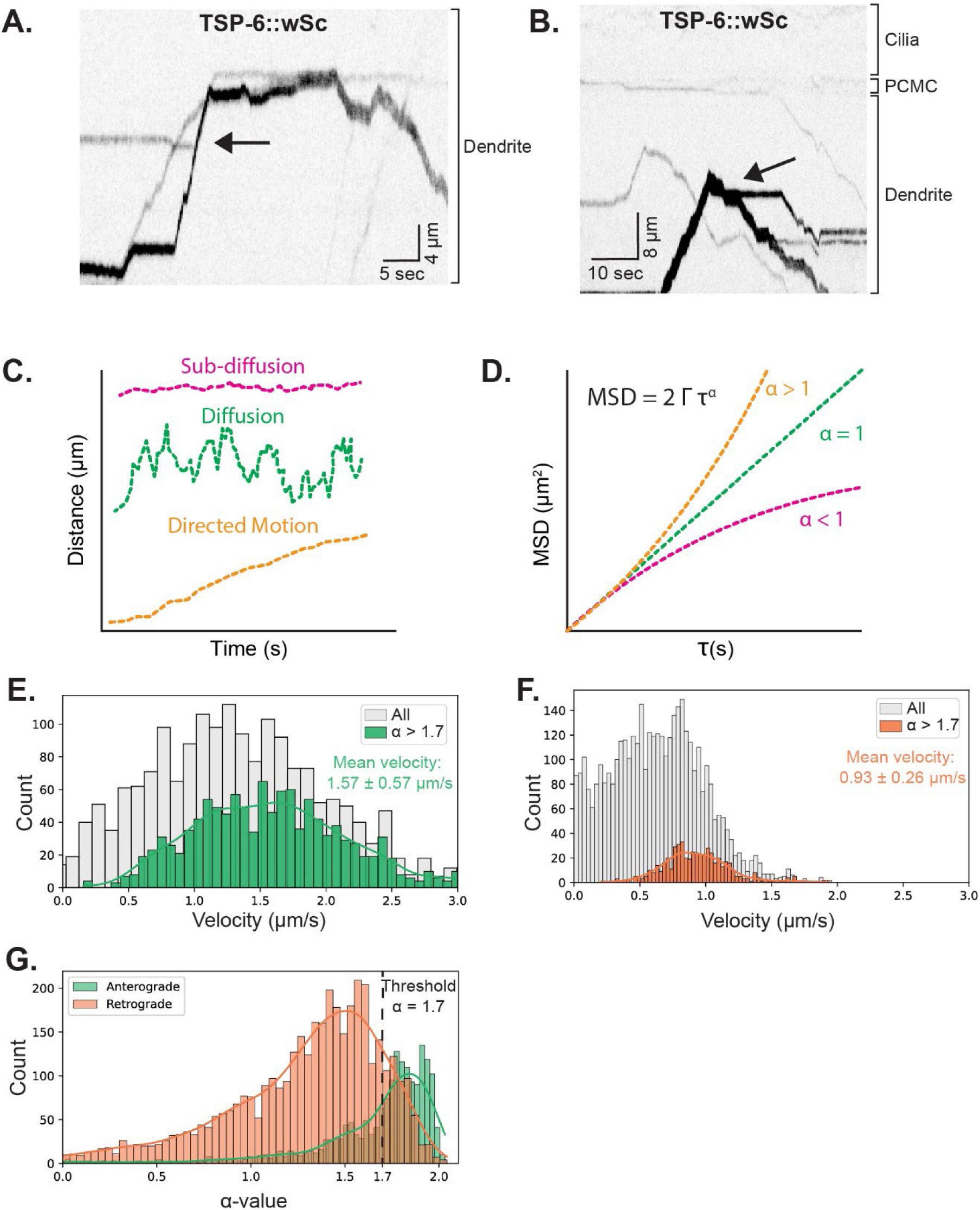
TSP-6::wSc dendritic vesicle velocity and α-value distribution and their mutual interaction. (A) Kymograph of TSP-6::wSc showing a fusion event. (B) Kymograph of TSP-6::wSc showing a fission event. (C) The plot shows distance along the cilium versus time with three example of particle motion, namely: sub-diffusion, diffusion and directed motion. (D) The plot shows the MSD curves associated with those three examples of particle motion. The anomalous exponent α is lower than 1 for sub-diffusive particles, 1 for diffusive particles and higher than 1 for particles with directed motion. (E-F) Histogram of TSP-6:wSc vesicle point-to-point velocity distribution in the anterograde (E) and retrograde (F) direction. In grey all data points are shown and in green and red the data points that are filtered with an α-value above 1.7. The green and red lines show the kernel density estimation. The mean and standard deviation of the filtered velocity are shown in the figure. (G) Histogram showing the point-to-point α -value distribution of all the anterograde and retrograde trajectories. The green and red lines show the kernel density estimation for the anterograde and retrograde distribution, respectively.

**Supplementary Figure 2:**
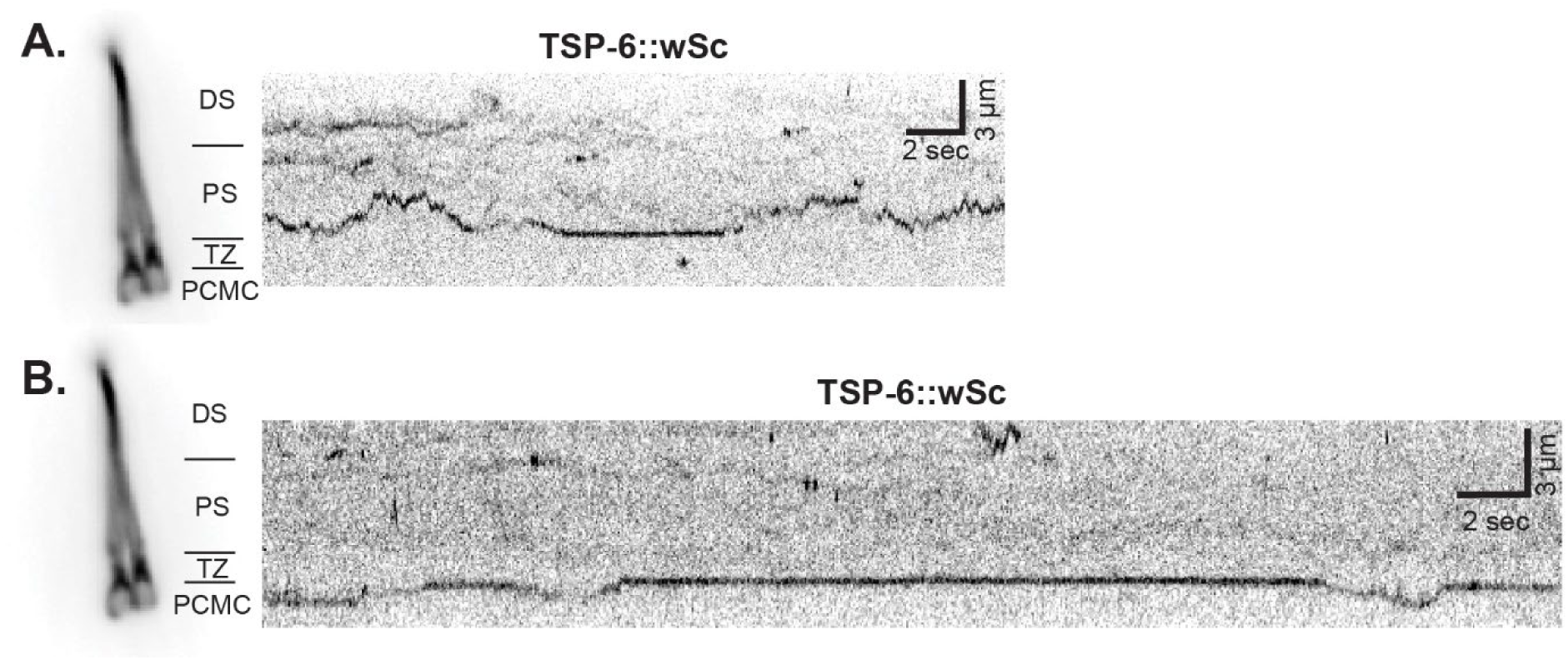
Transient changes between free diffusion and sub-diffusion in the transition zone. (A) Kymograph illustrating a single TSP-6::wSc molecule transitioning between free diffusion within the proximal segment and sub-diffusion within the transition zone, and returning to free diffusion within the proximal segment. (B) Kymograph showing a single TSP-6::wSc molecule transiently changing between free diffusion within the PCMC and sub-diffusion within the transition zone.

**Supplementary Figure 3:**
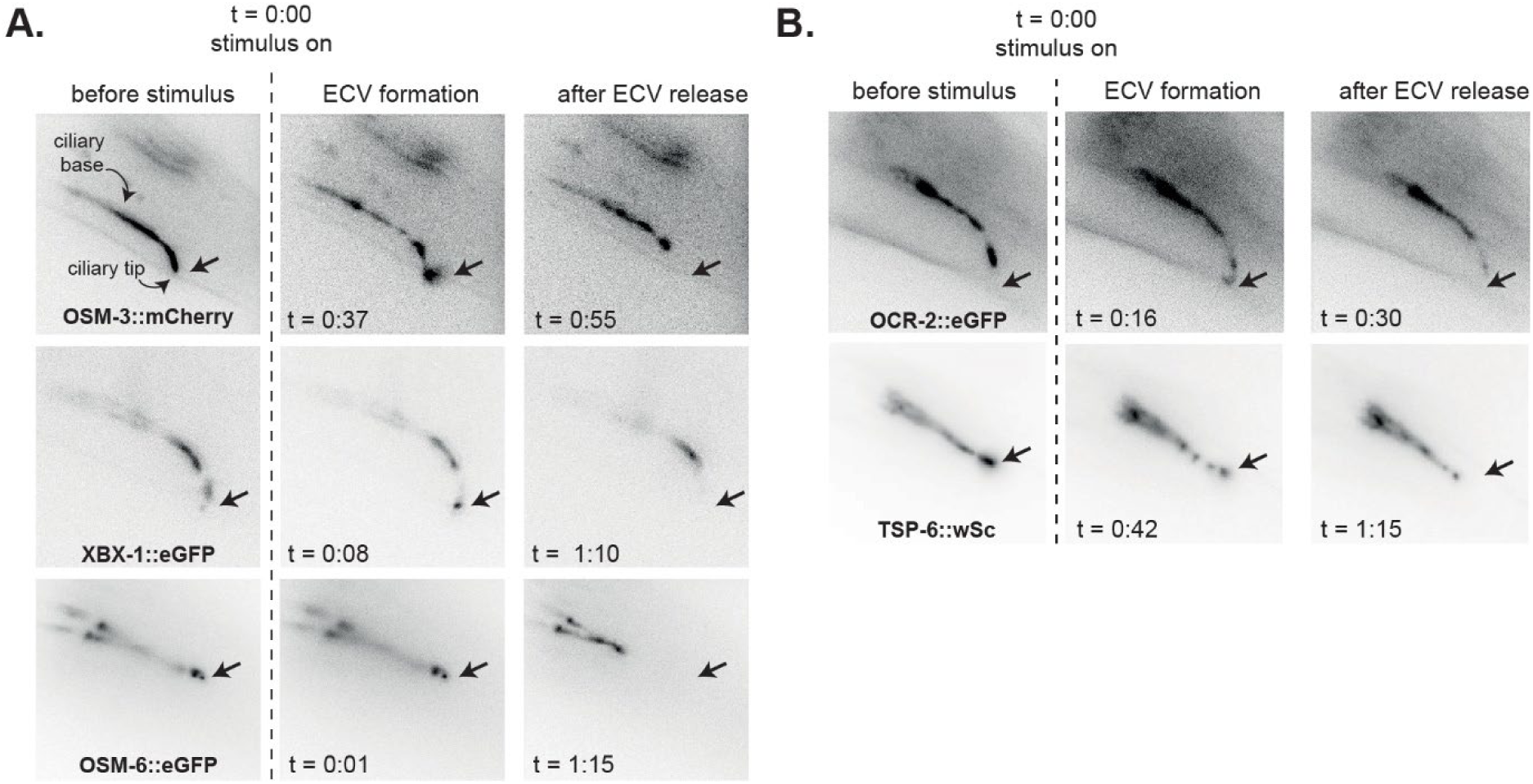
Various ciliary components are secreted by ectosomes. (A) Fluorescence images of OSM-3::mCherry (upper), XBX-1::eGFP (middle) and OSM-6::eGFP (lower), before 0.1% SDS exposure (left), during exposure to SDS, which leads to the shedding of material from the ciliary tip (middle), and after the release of the material into the environment (right). Position of ectosome release is indicated by a black arrow. (B) Inverted fluorescence images of OCR-2::eGFP; and TSP-6::wSc; before SDS exposure (left), during exposure to SDS, which leads to the shedding of material from the ciliary tip (middle), and after the release of the material into the environment (right). Position of package release is indicated by a red arrow.

**Supplementary Figure 4:**
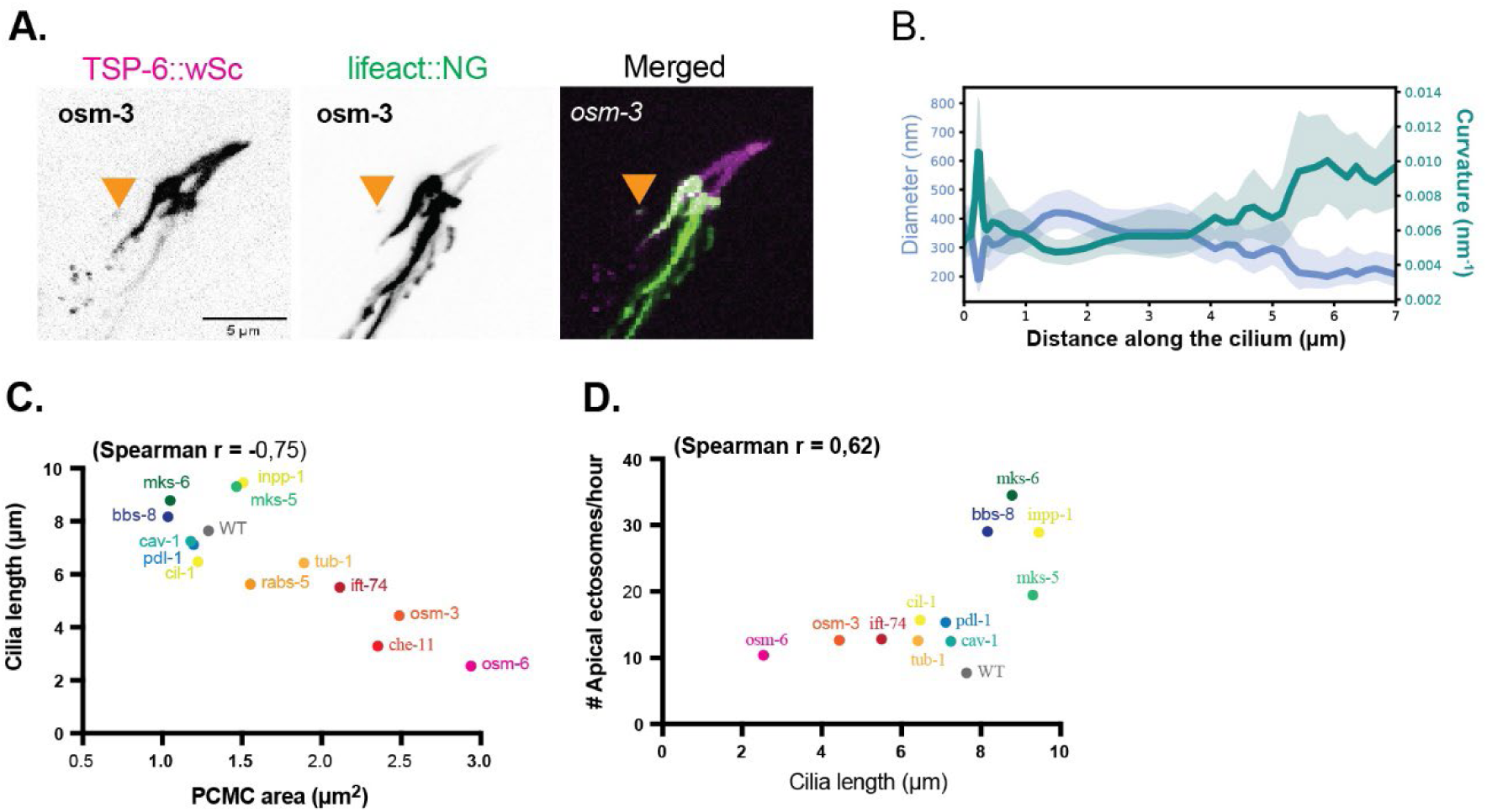
correlations between cilia morphologies and ectocytosis. (A) In *osm-3*, Lifeact::NG strongly stain the PCMC and distal dendrite but rarely enter basal ectosome (orange arrow). (B) Diameter and curvature along the cilium. The diameter is estimated by taking the 80^th^ percentile of the cumulative distribution of perpendicular distances derived from single-molecule localizations. The curvature is calculated as the inverse of the radius. The shaded area indicates the 70^th^-90^th^ percentile and indicates the error. (C-D) Correlation between cilia morphology parameters (cilia length and PCMC area) and ectocytosis location (basal or apical). Spearman rank suggests strong anticorrelation between cilia length and PCMC area (C) and a weak correlation between between cilia length and apical ectocytosis rate (D).

## SUPPLEMENTARY VIDEOS

**Videos S1-S6 – Colocalization of TSP-6::wSc with other organelles:** The videos present series of z-stack images, with each frame corresponding to a single z-step. Arrows highlight spots where colocalization occurs. TSP-6::wSc is represented in magenta, the tested markers labelled with StG/GFP are represented in green. Video S1 – Colocalization of TSP-6::wSc with StG::RAB-8 involved in trafficking from trans-Golgi. Video S2 – Colocalization of TSP-6::wSc with the early endosome marker StG-RAB-5. Video S3 – Colocalization of TSP-6::wSc with the recycling endosome marker StG::RAB-1. Video S4 – Colocalization of TSP-6::wSc with the lysosome marker LMP-1::GFP. Video S5 – No colocalization was observed between TSP-6::wSc and late endosome marker StG::RAB-7. Video S6 – Colocalization of TSP-6::wSc with the recycling endosome marker StG::RAB-11 within ciliated ending.

**Video S7-S8 - Dendritic TSP-6::wSc vesicle arriving at the PCMC:** Representative movie of a TSP-6::wSc vesicle moving in the dendrite towards the cilia, briefly remaining stationary in the PCMC before moving back in the opposite direction. The TSP-6::wSc fluorescence in the cilia has been photobleached away prior to this movie for better visualization of vesicle interaction with the PCMC, therefore the location of the PCMC and cilia are indicated. The arrow points to moment where the vesicle might be partly detaching before it moves back towards the soma. Video speed is about 4.5X. (S8: Same data as in Figure 1I.)

**Video S9 - Dendritic TSP-6::wSc vesicle arriving at the PCMC:** Representative movie of a TSP-6::wSc vesicle moving in the dendrite towards the cilia, remaining stationary while the fluorescent signal decays. The TSP-6::wSc fluorescence in the cilia has been photobleached away prior to this movie for better visualization of vesicle interaction with the PCMC, therefore the location of the PCMC and cilia are indicated. Video speed is about 4X. (Same data as in Figure 1J.)

**Video S10 - Response of TSP-6::wSc to acute copper stimulus:** Representative movie of TSP-6::wSc in response to exposure to 10 mM CuSO_4_ for 30 seconds. The start of exposure starts at t = 0:30 and ends at t = 1:00, as indicated in the video. Video speed is about 10X.

**Video S11 - Apical ectosome release after acute SDS stimulus:** Representative movie of the IFT machinery, in this case OSM-3::mCherry, in response to exposure to 0.1% SDS. The start of exposure starts at t = 0:15 and does not end during this movie. During exposure multiple apical ectosomes are released from the tip. Video speed is about 7.5X. (Same data as in Figure 3C.)

**Video S12 - Basal ectosome release after acute copper stimulus:** Two-color movie of TBB-4::eGFP and TSP-6::wSc after exposure to 10 mM CuSO_4_. During the recovery phase after exposure to the stimulus a basal ectosome containing TSP-6::wSc is released (indicated with the white arrow). Video speed is about 150X. (Same data as in Figure 3D.)

**Video S13 – cEVs bud from the proximal region of the cilia in arl-13 mutants:** Time-lapse imaging of PHA/PHB neuron cilia of *arl-13* mutant suggests TSP-6::wSc budding occurs in the proximal segment of the cilia. The arrow in the video indicates the site of ectosome emergence, the bracket labeled “c” highlights the cilium, while the other bracket indicates PCMC.

**Video S14 – In *mks-6* mutants, a sequence of events leads to apical ectocytosis including extension of ciliary tip, pinching and ectosome release:** Time-lapse imaging of PHA/PHB neuron cilia in *mks-6* mutant captures ectocytosis dynamics. Arrows indicate the sites of ectocytosis, while the bracket labeled “c” highlights the cilium.

**Videos S15-S16. The shape of the PCMC is highly dynamic in *osm-3* mutants:** Video S15 - Time-lapse imaging of PHA/PHB neuron cilia in *osm-3* mutant demonstrates the dynamics of branches formation from PCMC. Arrow indicates the site of branch formation, while the bracket labelled “b” highlights an already formed branch. Video S16 – Time lapse imaging of PHA/PHB neuron cilia in *osm-*3 mutant demonstrates basal ectosome release from a transient branch. Arrow indicates the cEVs budding and release, the bracket labelled “c” highlights cilium, while the other bracket highlights PCMC. On both videos (S15-S16), TSP-6::wSc labelled cilia are represented in gray scale, whereas the glia is represented in cyan.

